# The mutational landscape of human olfactory G protein-coupled receptors

**DOI:** 10.1101/2020.05.29.121103

**Authors:** Ramón Cierco Jimenez, Nil Casajuana-Martin, Adrián García-Recio, Lidia Alcántara, Leonardo Pardo, Mercedes Campillo, Angel Gonzalez

**Affiliations:** Laboratori de Medicina Computacional, Unitat de Bioestadística, Facultat de Medicina, Universitat Autònoma de Barcelona, E-08193 Bellaterra, Spain

## Abstract

Olfactory receptors (ORs) constitute a large family of sensory proteins that enable us to recognize a wide range of chemical volatiles in the environment. By contrast to the extensive information about human olfactory thresholds for thousands of odorants, studies of the genetic influence on olfaction are limited to a few examples. Here, we analyzed a compendium of 118,057 natural variants in human ORs collected from the public domain. OR mutations were categorized depending on their genomic and protein contexts, as well as their frequency of occurrence in several human populations. Functional interpretation of the natural changes was estimated from the increasing knowledge of the structure and function of the G protein-coupled receptor (GPCR) family, to which ORs belong. Our analysis reveals an extraordinary diversity of natural variations in the olfactory gene repertoire between individuals and populations, with a significant number of changes occurring at structural conserved regions. A particular attention is paid to mutations in positions linked to the conserved GPCR activation mechanism that could imply phenotypic variation in the olfactory perception. An interactive web application (available at http://lmc.uab.cat/hORMdb) was developed for the management and visualization of this mutational dataset.

## INTRODUCTION

Vertebrate olfactory systems have evolved to sense volatile substances through their recognition by olfactory receptors (ORs) located on the membrane of olfactory sensory neurons in the olfactory epithelium [1] and consequent initiation of signaling cascades that transform odorant-receptor chemical interactions into electrochemical signals [2, 3]. These receptors belong to the class A G protein-coupled receptors (GPCRs), a major drug target protein family [4] involved in the transduction of extracellular signals through second messenger cascades controlled by different heterotrimeric guanine nucleotide-binding proteins (G_olf_ in the case of ORs) coupled at their intracellular regions [5, 6].

ORs are characterized by intronless coding regions of average length of 310 codons (∼1kb) and constitute the largest multigene family in humans, with a functional gene repertoire estimated between 322 and 390 genes, divided into two main classes, 17 families and more than 150 subfamilies [7]. This broad array of receptors, like in other terrestrial mammals, is shared with tetrapods (families 1-14) and marine vertebrates (families 51-56) [8], and seems necessary to respond efficiently to the extraordinary chemical diversity of odorants in Earth’s ecosystems [9]. However, there is growing evidence that their functional roles are beyond olfactory tissues [10, 11].

Human genomic data reveal that loci of human ORs harbor a considerable number of genetic variability [12]. Many of these variants may interfere with the receptor expression, interaction with odorants or signal transduction, and consequently could modify the physiological response to a determinate olfactory stimulus. In this regard, it has been long established a considerable variation in the perception of odorants among individuals [9, 13] and populations [14, 15], which in some cases has been associated to genetic changes in OR genes [16-18]. To further study this issue, we used publicly available human sequencing data to conduct *in-silico* data mining and analysis of ORs natural variants in 141,456 human exomes and genomes from more than one hundred thousand unrelated individuals [19].

Information of chromosomal localization, type of substitutions and allele frequencies in several sub-continental populations were obtained for close to a hundred and twenty thousand natural variants identified in 374 human ORs. A detailed topological localization system was developed to assign each mutation to a region within the seven alpha helical bundle molecular architecture characteristic of GPCRs (i.e. extracellular and intracellular N- and C-terminal sequences, seven transmembrane α-helices [TM 1 to 7], three extracellular [ECL 1 to 3] and three cytoplasmic loops [ICL 1 to 3]) [20, 21]. This system also includes the assignation of unambiguously positions to all mutations occurring in the TM helices according to the numbering systems developed by Ballesteros-Weinstein (BW) and others for this family of proteins [22, 23].

The analysis of the collected data revealed numerous differences among individuals and populations, with an allele frequency spectrum dominated by low frequency variants. A significant number of natural changes were identified at GPCR functional regions [24-28], or forming part of ligand binding cavities [29]. These, and the rest of the coding-sequence mutations were evaluated according to an amino acid substitution score weighting developed for this family of receptors [30]. The utility of this topological annotation approach is illustrated with selected examples of natural OR variations that could imply phenotypic changes in the odorant perception for a substantial group of individuals. These results are accompanied by a computational application developed to facilitate the public access and analysis of this data. The human Olfactory Receptor Mutation database (hORMdb, https://lmc.uab.cat/hORMdb) is an interactive database, that allows the selection and filtering of human OR natural variants order to analyze specific dbSNP entries, individual genes or complete families according to their topological localization, population frequencies and substitution scores, amongst other features.

## MATERIAL AND METHODS

### Data acquisition and filtering

Natural sequence variations from functionally annotated human ORs [31, 32] were obtained from the Genome Aggregation Database (gnomAD, http://gnomad.broadinstitute.org/) using Python (v.3.7.6) data mining scripts. Variant tables for each OR were imported to R (v.3.6.2), including information of chromosome location, transcript consequence and allele frequencies in seven sub-continental populations [19] (Supplementary Table S1). BLAST (v.2.10.0) and Python scripts were used to compare the collected sequence information with UniProt database (release 2019_11, https://www.uniprot.org/). The collected data was then filtered to remove null values, duplicates, missing rsIDs and sequence conflicts with reference Swiss-Prot entries, resulting in a curated dataset of 118,057 nucleotide variants from 374 human OR genes (Supplementary Table S2).

### Topological mapping and BW annotation

Python data mining scripts were used to assign each coding-sequence mutation a topological location according to a structure-based MSA of the human ORs Swiss-Prot reference sequences and class A GPCRs of known three-dimensional structure (Supplementary Figures S1 and S2). Natural variants at the TM regions were further annotated with the generic two number system developed by BW consisting of two digits: the first (1 through 7) corresponds to the helix in which the change is located, and the second indicates its position relative to the most conserved residue in the helix (arbitrarily assigned to 50) [22]. This nomenclature was also applied to a 10 residues stretch located between two highly conserved cysteines at the ECL2 (indicated by 45 as first number attending to its location between the TMs 4 and 5) [23].

### Impact evaluation of coding sequence variants

Impact of non-synonymous changes was estimated from the amino acid substitution scores derived from the GPCRtm matrix [30] (Supplementary Figure S5). In addition, two subsets of BW topological sites were outlined: *i)* a functional core (FC) subset of 30 topological positions with high degree of conservation likely involved in the receptor activation, G-protein binding or disulfide bond formation (Supplementary Table S3 and Figure S3); and *ii)* a binding site (BS) subset of 30 amino acid positions within a distance of ≤ 4.0 Å to bound ligands in 39 reference class A GPCR 3D structures (Supplementary Table S4 and Figure S4).

### Development of an interactive database with the annotated variation data

Substitution scores and topological annotation (including FC/BS and BW numbering) were transferred to the mutation data table using Python data mining scripts, completing the annotation process (Figure 1 and Supplementary Table S5). A standalone application was programed with the open source RStudio (v.1.2.5003) to manage and visualize this curated mutation dataset (https://github.com/lmc-uab/hORMdb). This database resource is also made available online as an interactive web server programed with the Shiny Server package (v.1.5.12.933) (http://lmc.uab.cat/hORMdb).

**Figure 1.**
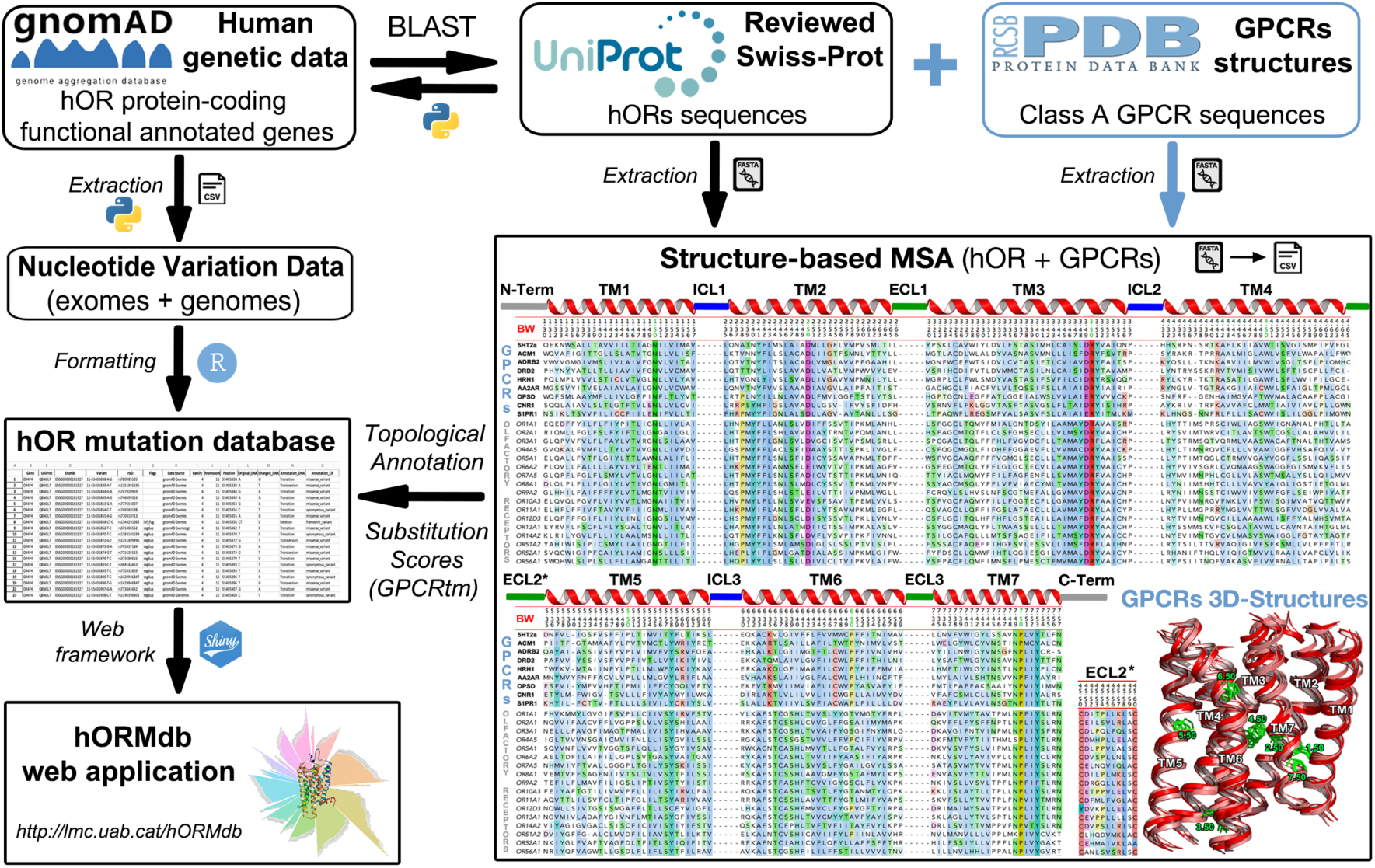
Data flow describing the extraction and topological annotation of OR natural variants. Functional human OR genes were used as queries for genotype searches in gnomAD and the results were stored in a mutation data table (left side of the diagram). BLAST searches were used to localize the corresponding OR UniProt sequences. Topological regions and BW notation were defined from a structure-based MSA with the ORs sequences and class A GPCRs with solved 3D atomic coordinates (right side of the diagram). This information and their associated substitution scores were transferred to each entry in the mutation data table, thus completing the annotation process. Full length sequences and accession numbers of the sequences used in the study are available at Supplementary Figures S1 and S2.

## RESULTS

Natural variations in human ORs were mined from nucleotide sequence data of 138,632 unrelated individuals in the Genome Aggregation Database [19], and annotated at structural level with information of the class A GPCR family as resumed in Figure 1. This curated dataset comprises 118,057 nucleotide changes in 374 functional OR genes, which belong to 17 OR families (Figure 2). Overall average number of mutations per receptor was 316, with a prominent variation rate in the OR52 family (average of 343) and five members of the OR4 family (OR4A5, OR4A15, OR4A16, OR4C16 and OR4C46) with more than 500 mutations/receptor. On the other hand, lowest variation rates correspond to the OR14 family (average of 256) and few more than a dozen receptors with less than 100 mutation counts (Supplementary Table S2). This staggered mutational distribution supports a heterogeneous selective pressure in OR genes, as indicated by other studies [33, 34].

**Figure 2.**
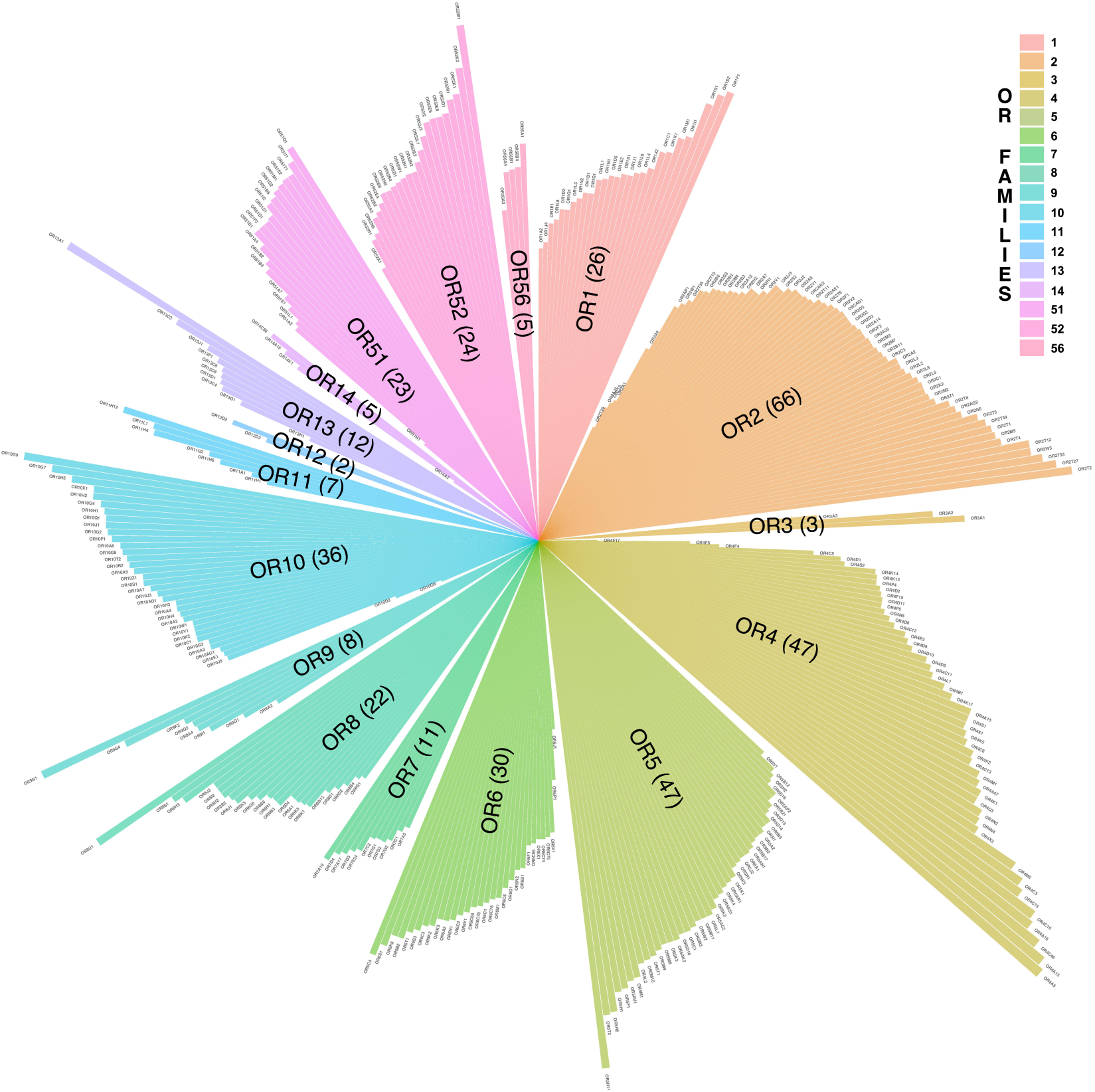
Wind rose plot representing the mutational landscape of human ORs. The plot shows the distribution of 118,057 nucleotide variants in 374 functional OR genes (color bars) clustered in 17 families (top left legend, ordered clockwise). Numbers in parenthesis correspond to the total number of receptors analyzed at each family. Gene names and family assignation correspond to the recommended terms by the HUGO Gene Nomenclature Committee (HGNC). Names of individual OR family members are in the format “ORnXm”: a root name ‘OR’, followed by family numeral (n), subfamily letter (X), and a numeral (m) representing the particular gene within the subfamily. For example, OR2A1 is the first OR gene in the family 2, subfamily A.

Most common variations types in the collected dataset correspond to missense (∼64%) and synonymous substitutions (∼25%), followed by frameshifts, non-coding (3′UTR and 5′ UTR), stop gained, and a reduced number of other minor mutations events (Figure 3A). Regarding the nature of the changes, transitions and transversions are the most likely mutational events, representing 96% of the entire dataset, whilst the remaining 4% correspond to deletions and insertions (inset on Figure 3A). The large OR multigene family occupies vast amounts of genomic territory. As expected, the number of mutations per chromosome is linked to the genome distribution of the OR genes (Figure 3B). Chromosome 11, which contains the largest number of receptors, displays the highest number of variants, followed by chromosome 1. For the rest of chromosomes hosting OR genes the number of variants ranges from ∼7,000 to less than 100, and no data was recorded for chromosomes 4, 8, 13, 18, 20, 21 and Y. A graphical display of the unevenly chromosomal distribution of the mutations in the 17 human OR families is available in the Supplementary Figure S6.

**Figure 3.**
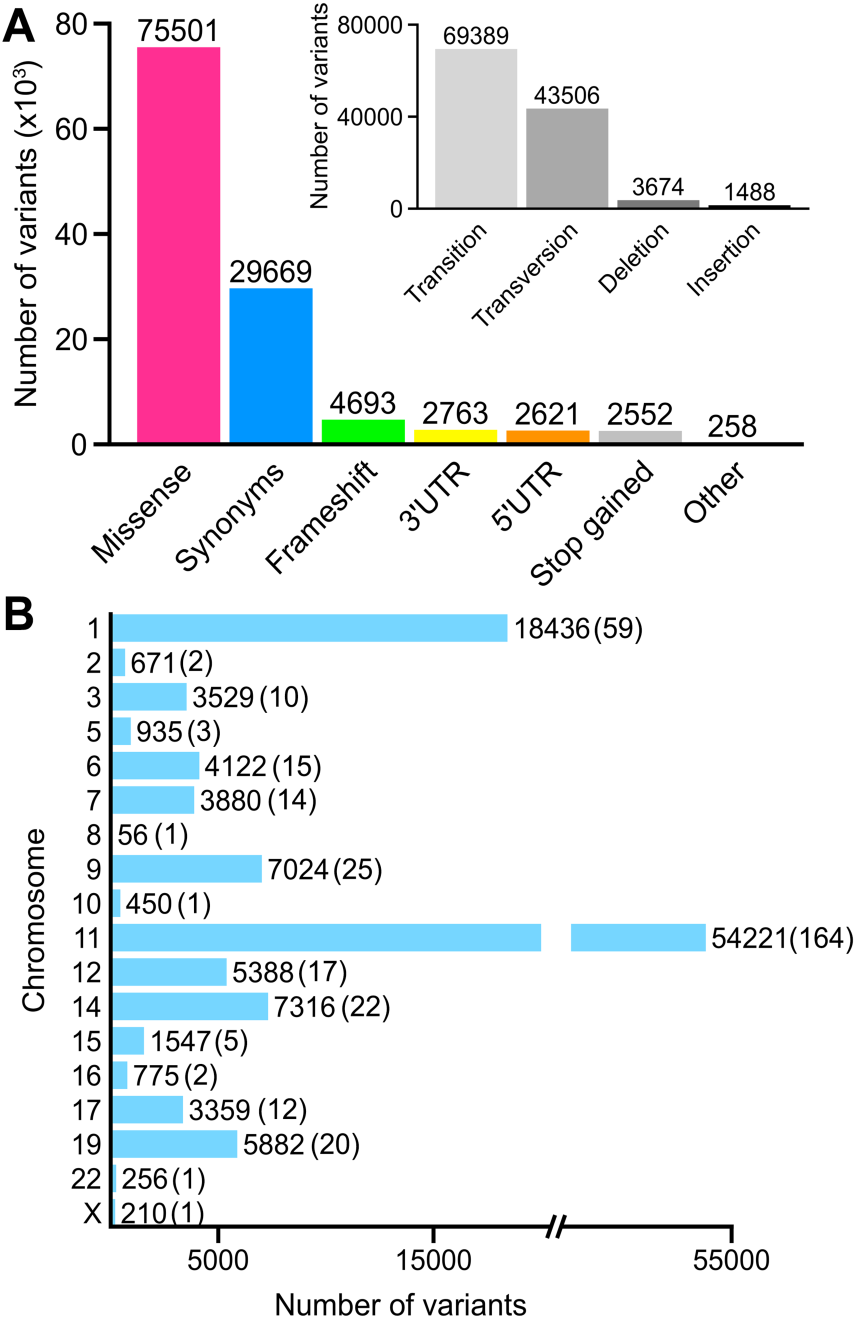
Categorization and chromosomal distribution of human OR natural variants. A. Functional categories and type of changes (inset) of the sequence variants identified in the 374 investigated OR genes. B. Number and distribution of the OR variants per chromosome (the number of OR analyzed per chromosome are shown in parenthesis).

### Allele frequencies and population distribution of the variant dataset

Analysis of the frequency values from gnomAD disclose only 2,152 OR natural variants with global allele frequency above 1% in the collected dataset. By contrast, >95% corresponds to low frequency variants (59,829 of which are singletons), exposing an extraordinary interindividual variation in the human OR gene repertoire. Taking into account that differences in olfactory sensitivity could be at least partly explained by the prevalence of particular mutated ORs alleles in individuals within populations [35]; independent frequency ranges were analyzed on each of the seven sub-continental populations in the database (Figure 4 A-G and Supplementary Table S1). This analysis shows a similar trend of frequency distribution among ethnic groups, characterized by an elevated number of mutations with allele frequencies below 0.1%. From these changes, 36,749 were exclusively found in the European (non-Finnish), 14,627 in South Asian, 11,095 in African, 10,487 in Latino, 9,855 in East Asian, 1,771 in Finnish and 811 in Ashkenazi Jewish populations.

**Figure 4.**
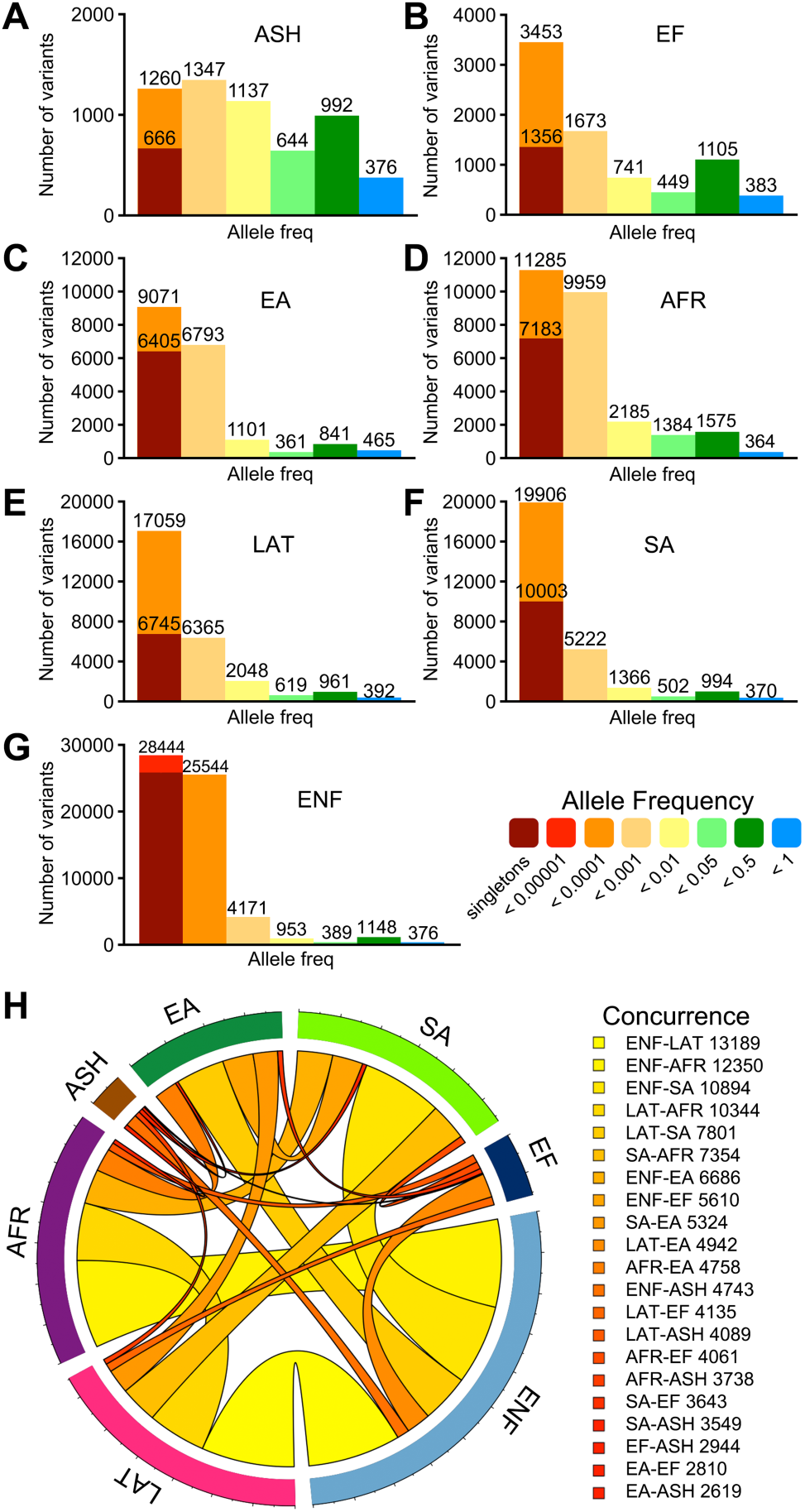
Allele frequencies and concurrence of OR mutations within human populations. Number and frequency distribution of natural variants from seven sub-continental populations obtained from gnomAD and corresponding to: A. Ashkenazi Jewish (ASH), B. European Finnish (EF), C. East Asian (EA), D. African (AFR), E. Latino (LAT), F. South Asian (SA) and, G European non-Finnish (ENF). Bars are colored according to the allele frequency scale on the right. H. Circos plot of the concurrence of mutations between populations. The width of the link between two populations corresponds to the number of shared mutations between them, also represented by a color gradient (numerical values are displayed on the right).

On another note, assessment of the concurrence of mutations reveal 2,097 genetic variants common to all populations, of which 1,812 display allele frequencies >1%. Notwithstanding, 28,880 variants were identified in two or more ethnic groups. Pair-wise comparisons of shared mutations between sub-continental populations are summarized in the circos plot of Figure 4H. As observed in the graph, the largest European (non-Finnish) population shares more variants with the rest of ethnicities. This data, expressed as percentage of the total number of mutations at each population indicates that approximately 82% of the Ashkenazi Jewish, 72% of Finnish, 48% of Latino, 46% of African, 39% of South Asian and 36% of the East Asian natural variants are shared with the European (non-Finnish) population. Likewise, African and Latino share ∼38% of mutations, whereas the South Asian population shares ∼27% of mutations with Latino, ∼26% with African, and surprisingly less than 20% with the East Asian population.

### Topological assignment to sequence variants

Topological domain assignation of coding sequence variants according to the conserved class A GPCR molecular architecture (i.e. N-term, 7-TMs, 3-ECLs, 3-ICLs and C-term defined in the structure-based MSA in Figure 1), revealed that 65% of the mutations were located in the TM regions (50,703 missense, 20,296 synonymous, 3,170 frameshifts and 1,657 stop gained variants) (Figure 5). TM6 accumulates more changes (13,117 variants), followed by TM3 (12,782) and ECL2 (11,057). On the other hand, a lower number of mutations were found in intracellular and extracellular loops, N- and C-terminal domains and NCRs. This trend is observed in all OR families, with most changes occurring in the TMs and ECL2, and major inter-family differences in the NCRs and N- and C-terminal domains because of their variable lengths (Supplementary Figure S7).

**Figure 5.**
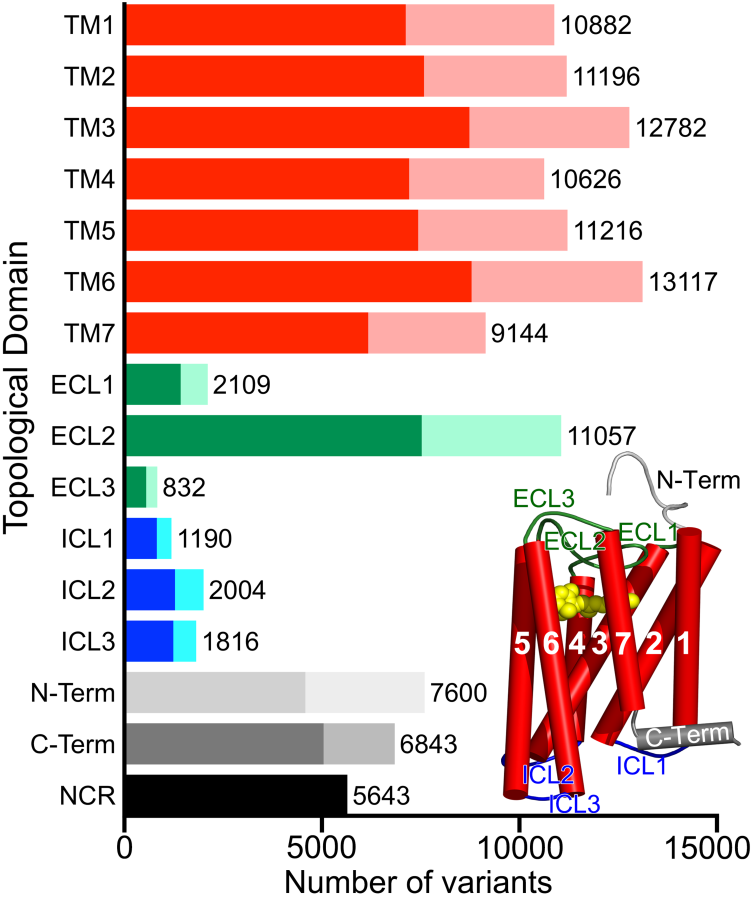
Number and distribution of mutations within topological domains. Number of variants (x-axis) per topological domain (y-axis) as defined by the conserved class A GPCR molecular architecture of seven TM helices (TM1 to 7 in red), three extracellular loops (ECL 1 to 3 in green), three intracellular loops (ICL 1 to 3 in blue) and N- and C-terminal regions (in grey). NCR stands for non-coding regions. Darkest regions on the y-axis bars correspond to missense substitutions. The cartoon model in the lower right exemplifies the extent and arrangement of the topological regions according to the crystallographic structure of Rhodopsin (PDBid: 4J4Q, retinal is shown in yellow VdW spheres).

The analysis of individual positions within the conserved GPCR topological domains, using the BW nomenclature, reveals that, overall, the occurrence of natural variants is not restricted to a specific TM region or particular location, with an average of 360 changes per site (Figure 6). However, position 3.50 (995 total variations, 816 missense) stand out from the rest of sites (Figure 6C). This conserved position constitutes a switch for the signal transmission mechanism, which involve the structural rearrangement of the TM regions, opening the intracellular cavity for G protein binding, through changes in the DR^3.50^Y interaction environment [36, 37]. Consequently, this position is very sensitive to natural sequence variations linked to pathological outcomes in several GPCRs [38-42]. Interestingly, most frequent substitutions of the conserved Arg^3.50^ (96% conservation in GPCRs, 92% in ORs) involved the amino acids His (192 occurrences) and Cys (185 occurrences), which is consistent with a previous study conducted on non-olfactory GPCRs [43].

**Figure 6.**
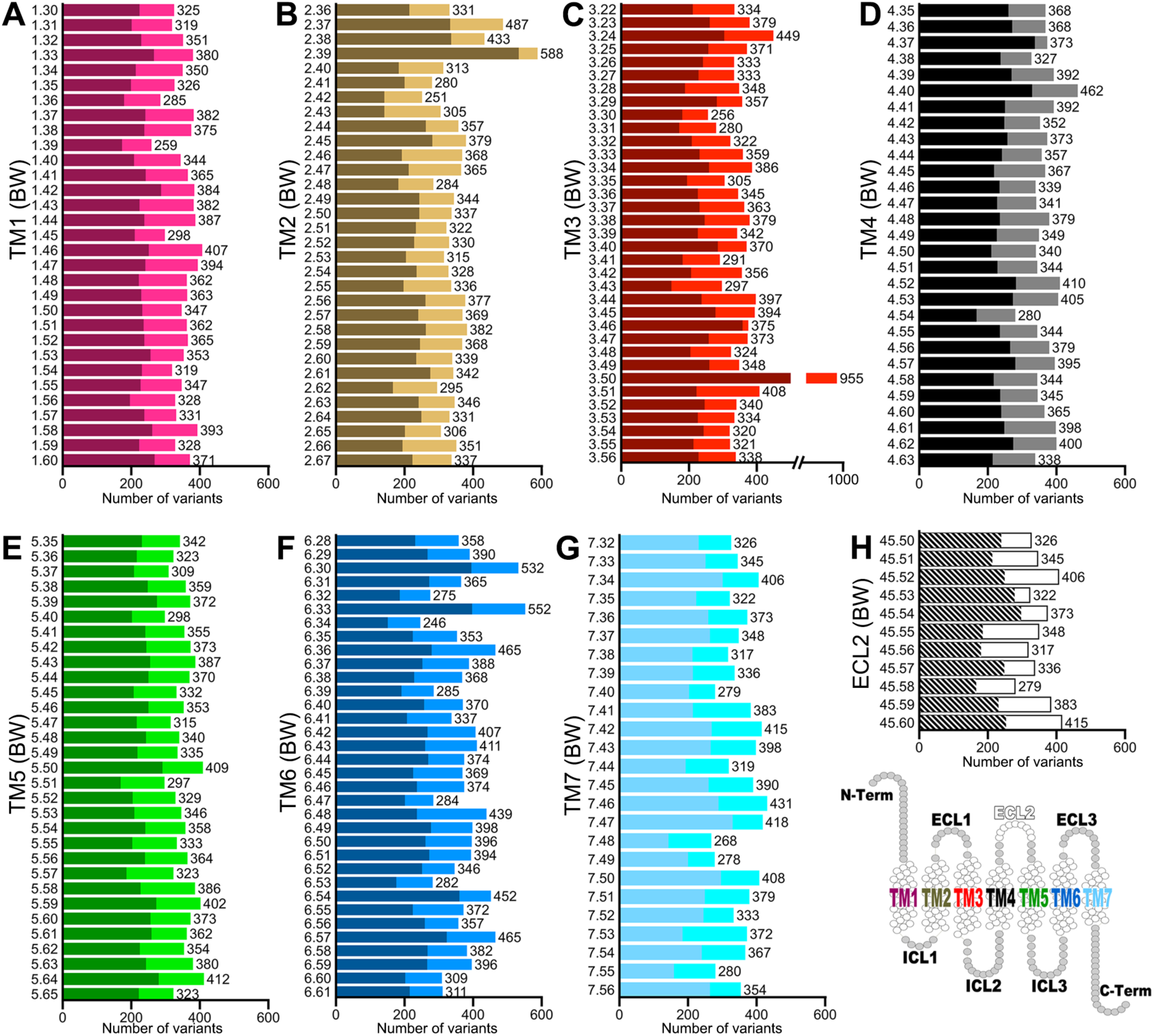
Number and distribution of mutations within the TM regions. Mutation counts (x-axis) associated to conserved topological sites at the seven-transmembrane (7-TM) and ECL2 regions (A to H). Positions of natural variants (y-axis) were assigned according to the Ballesteros-Weinstein (BW) numbering system derived from their respective positions on the structure-based MSA (see methods). Darkest regions on the y-axis bars correspond to missense substitutions. Empty circles in the snake-plot at bottom-right indicate the topological positions analyzed.

### Mutability landscapes of amino acids changes

Single amino acid variants in human ORs can alter the resulting phenotype, for example, by altering the odorant perception [44]. Thus, we investigate the type and magnitude of the amino acid changes in missense substitutions (75,501 variants in the dataset) as a first approximation to evaluate their functional consequences at the molecular level. As displayed in Figure 7A, hydrophobic residues (Leu, Ile, Val and Ala) exhibited the highest levels of mutability, followed by Ser and Thr in agreement with their stabilization roles on the structure of TM helices [45, 46]. Conversely, substitutions of Trp or polar/charged Gln, Glu, Lys, Asp, His and Asn (often associated with protein malfunction in TM proteins) were less frequent [47, 48].

**Figure 7.**
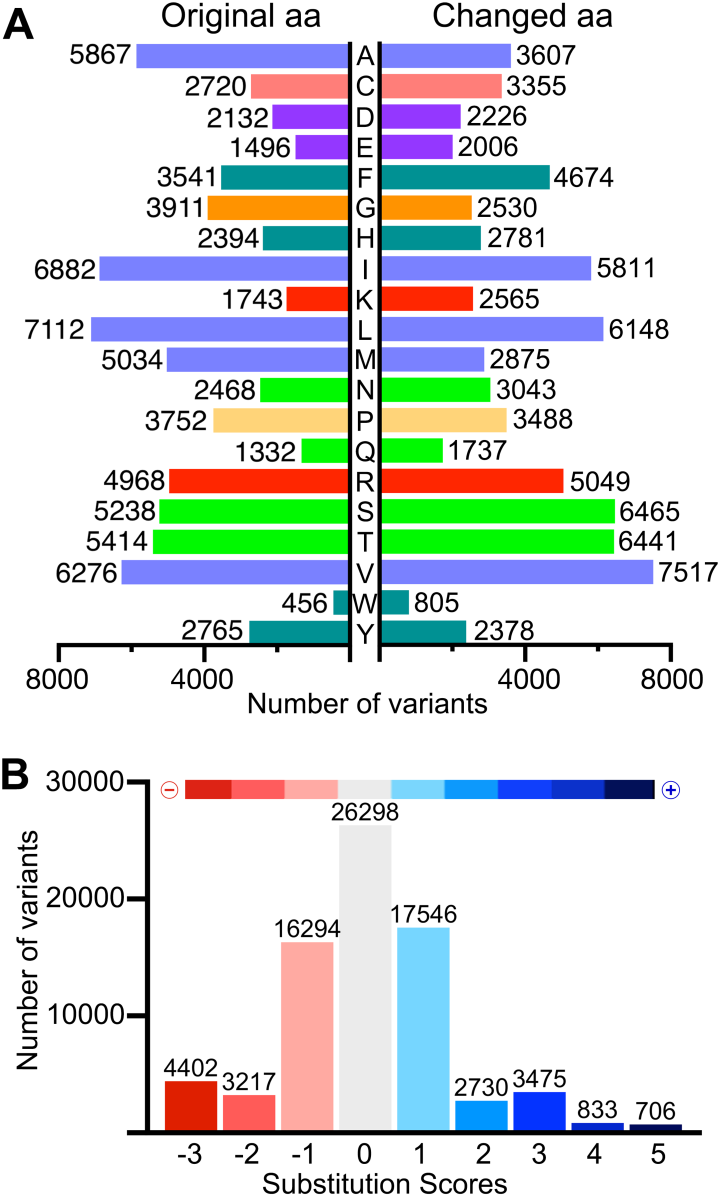
Mutability landscapes of amino acid substitutions in human ORs. A. Number of original (left) vs. changed (right) amino acids due to missense substitutions. Amino acid bars are colored by physico-chemical properties of the residues (Hydrophobic=blue, green=polar, cyan=aromatic, purple=negatively charged, red=positively charged, special residues Cys, Pro and Gly in yellow). B. Categorization of all amino acid replacements according to substitution scores extracted from GPCRtm. The color scale bar on top indicate the range of score values for all computed changes (negative=red, positive=blue).

The evaluation of the magnitude of changes was conducted using amino acid substitution scores derived from more than one thousand class A GPCR sequences (including ORs) and thus, reflecting the compositional bias distinctive of this particular family of proteins [30] (Figure 7B and Supplementary Figure S5). From this analysis, ∼75% of the missense substitutions were associated to zero or positive substitution scores (57,588 variants), indicating a preservation of physico-chemical properties of the original residue. Nonetheless, 24,561 changes compute negative scores, reflecting significant differences between the original and substituted amino acid, with possible impact on the receptor structural integrity and/or the binding of odorant molecules.

### Use of topological annotation, substitution metrics and allele frequencies in the impact evaluation of the mutations

Topological mapping of natural variations and their associated substitution scores were used in the functional imputation of missense substitutions. These features were analyzed in two subsets of topological positions within the conserved TMs and ECL2, which could either be involved in the receptor integrity and functional mechanism (Functional core, FC) or, in odorant-receptor interactions (Binding Site, BS) (Supplementary Figures S3 and S4). FC and BS topological subsets comprise 60 BW annotated positions and accumulate 7,437 and 7,330 missense variants counts respectively, of which 5,507 computed negative substitution scores. From this data, we identify 80 changes with allele frequencies > 1% in at least one of the sub-continental populations that could implicate distinctive odorant sensitivities for a considerable group of carriers (Supplementary Figure S8). At the moment, based on the limited published information of known ligands for human ORs we can only hypothesize about the impact of such changes through a few concrete examples described below:

#### Extracellular loop 2 at the conserved Cys45.50

A conserved cysteine residue in this position is involved in a disulfide bridge between ECL2 and TM3 in > 80% of class A GPCRs, and its substitution is related to a loss of function [28, 49, 50] (Figure 8A, B and E). An example of this type of mutation is found in the OR8B4, a recently deorphanized receptor for anisic aldehyde and muguet alcohol [51]. Variation rs4057749 (c.532T>C, p.Cys178Arg) in the OR8B4 may lead to impairment in the ability of perceive these aromatic cosmetic substances in a considerable proportion of the population (Supplementary Figure S8).

**Figure 8.**
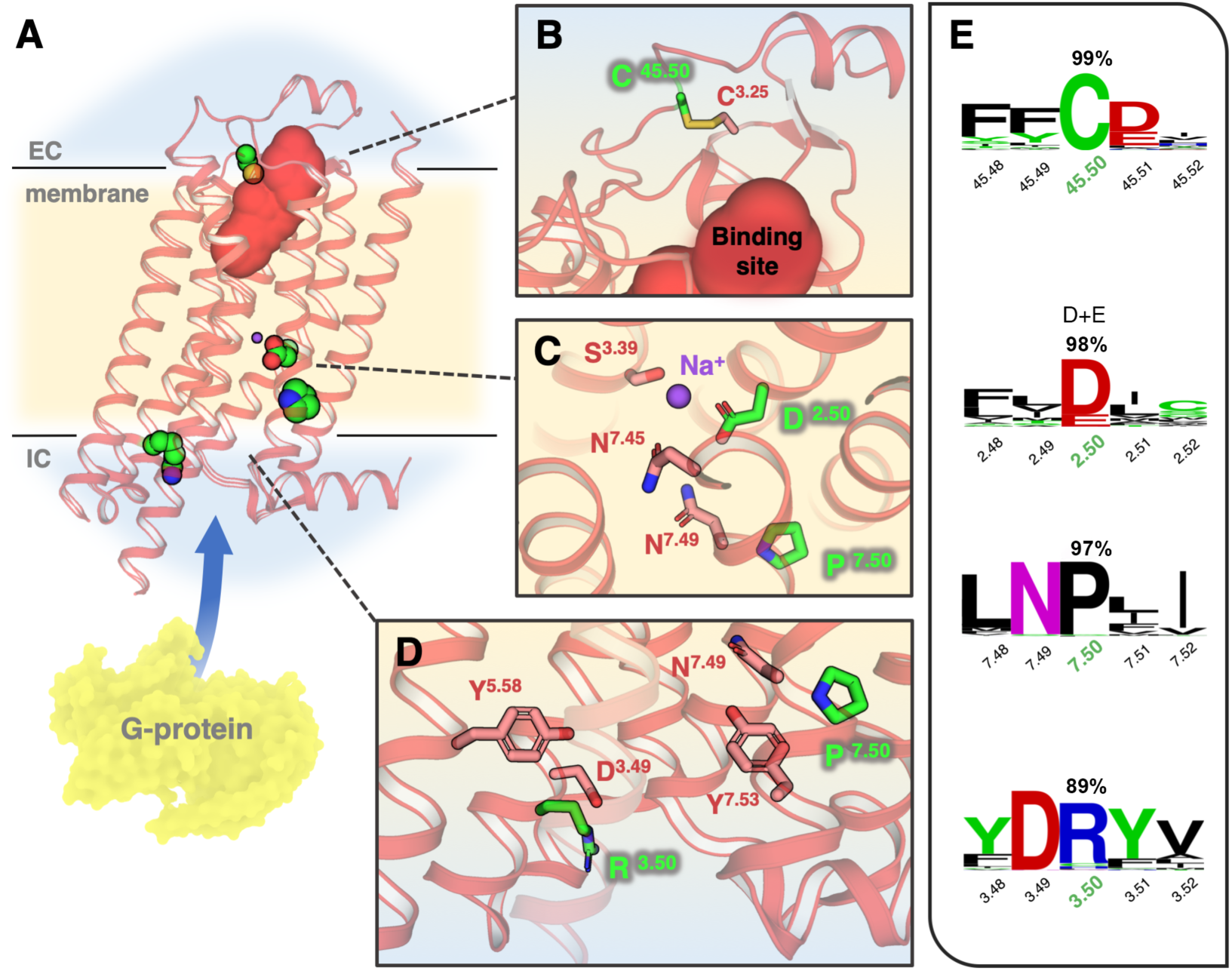
Structural visualization and examples of paradigmatic mutations selected from the study. A. General view of the adenosine A2A receptor (PDBid: 4EIY) as a prototype GPCR with the TM boundaries indicated in light yellow and ECLs/ICLs regions in blue. The ligand binding site is indicated by a solid red surface and G-protein interacting site by a blue arrow. Selected topological positions C^45.50^, D^2.50^, R^3.50^ and P^7.50^ are highlighted in the structure as green vdW spheres. B-D. Closer look of the atomic environment of selected positions (green sticks) and surrounding residues (salmon). E. Sequence conservation logos around the selected positions (number corresponds to their conservation percentage in the MSA). All the residue positions are referenced following the BW convention.

#### Transmembrane helix 2 at the conserved Asp2.50

It is characterized by the presence of a negative ionizable residue in the conserved (N/S)LxxxD^2.50^ motif, which is involved in the GPCR activation mechanism through allosteric modulation mediated by ionic species [52] (Figure 8A, C and E). Replacement of the conserved D^2.50^ would impair the coordination of modulating ions due to the loss of the negatively ionizable center [53]. Carriers of mutations on this site, as the rs4501959 (c.262G>A, p.Asp88Asn) in the OR52L1 might have different abilities to perceive carboxylic acids present in human sweat [54] and some of the components from the butter smell like butanoic acid and gamma decalactone that interact with this receptor [55].

#### Transmembrane helix 7 at the conserved Pro7.50

A conserved Pro in this position forms part of the NP^7.50^xxY motif involved in the transition from the ground state to the active forms of the GPCRs and internalization [56] (Figure 8A, C, D and E). Substitution of the P^7.50^ would modify the TM7 conformation producing a change of signalization patterns as observed in Rhodopsin [27]. An example of mutation on this site is found in the OR1A1, rs769427 (c.853C>T, p.Pro285Ser), which probably would affect their carriers for the detection of citronellic terpenoid substances identified as ligands for this receptor [57].

#### Transmembrane helix 3 at the conserved Arg3.50

A conserved Arg is the central component in the DR^3.50^Y motif directly implicated in the general activation mechanism of the class A GPCRs and its substitution generally modifies the transduction capacity of the receptor [24, 38-42] (Figure 8A,D and E). Natural variations at this position are found in most ORs, some of them at moderate to high frequencies in the populations investigated; examples include rs2072164 in OR2F1, rs3751484 in OR6J1, rs10176036 in OR6B2, rs12224086 in OR5AS1, rs2512219 in OR8D2, rs16930982 in OR51I1 and rs11230983 in OR5D13.

### Development of an interactive application to explore the human OR mutation data

It is expected that progress on OR genome association studies will continue to be made in the future. Thus, an interactive computational application was developed for the free access and analysis of this data by academics and industry professionals. The human Olfactory Receptor Mutation database (hORMdb, http://lmc.uab.cat/hORMdb) provides a curated and downloadable repository of natural variations in human ORs and several interactive tools for the selection, filtering and analysis of its contents (Figure 9).

**Figure 9.**
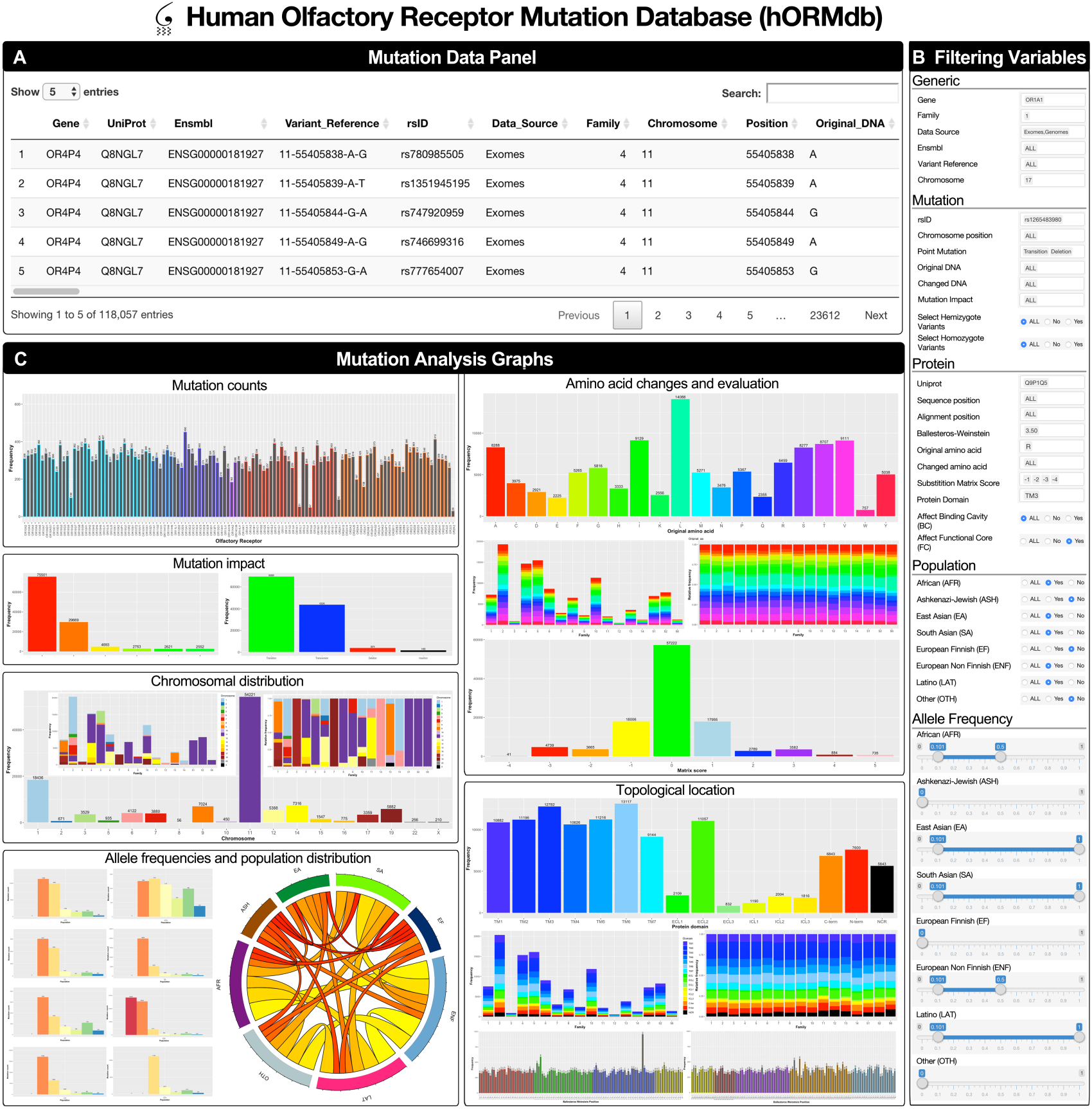
Overview of the hORMdb web interface. The hORMdb is an online resource to study natural variations in human ORs and it is structured in three main panels. A. A mutation data panel containing a downloadable table of the natural variants with multiple columns of information. B. A filtering variable panel allow the selection, concatenation and filtering of the data. C. A graphical panel for the visualization of the mutation data through several interactive graphs. Further details of the contents and organization of the database are provided in the HELP panel at the online application (http://lmc.uab.cat/hORMdb).

The hORMdb is structured as a data table (Figure 9A), containing information about individual dbSNP entries, particular genes or entire OR families, including the types of nucleotide and amino acids changes, allele frequencies in several sub-continental populations [58], and topological location in the receptor structure. All the mutation data can be selectively accessed through a filtering variable panel (Figure 9B) that allows the possibility of concatenate multiple selection choices (including numerical ranges for allele frequencies) or predefined topological subsets to analyze (e.g. FC, BS). Finally, a graphical panel interface (Figure 9C) allows to interactively display the selected content according to receptor types, chromosomal location, mutation impact, original/changed amino acids, substitution score, topological domain, BW position, allele frequencies and concurrence within populations. Altogether, this tool is intended to be used for the functional assessment of natural variations, rationalization of mutation data experiments or for comparative population studies.

## DISCUSSION

It is common ground that olfactory sensitivity differs across individuals and, in some cases, this feature has been related to genetic variations. Thus, the contribution of the genotype in the perception of odorants and volatile chemical mixtures seems particularly relevant. The highly diverse ORs, at the membrane of the olfactory neurons, trigger the first input of the olfactory signal. Thus, genomic studies of this family of receptors represent an important source of knowledge for academics and industry professionals who study the human olfaction. To this end, we can take advantage of the vast amount of information of natural genetic variations coming from genome-data community shared initiatives freely available in the public domain.

Using data mining tools, close to 120,000 nucleotide variations in human ORs were obtained from the large-scale sequencing data repository gnomAD, that provides well-structured information of sequencing data from a wide variety of sequencing projects all over the world [59]. The curation and computer analysis of this variation data revealed an uneven distribution of mutations in OR genes, reflecting the active role of natural selection in this family of receptors. Moreover, a considerable proportion of the identified mutations occur at very low frequencies, many of them uniquely identified at definite ethnic groups or individuals. This extraordinary genotypic variation has been earlier described [60], and suggest a great phenotypic diversity in the olfactory perception between humans.

Taking into account the increasing need of tools providing accurate predictions of functional consequences of natural variants identified in genomic studies [61]; evolutionary conservation and structural context were considered as key elements in the estimation of functional role of the natural variations identified. It is worth stressing that, in many cases, the structural framework of the mutated sites (intimately linked to the stability, function and interactions) is often overlooked due to a limited structural knowledge [62]. ORs are not an exception to this reality, with no molecular structure reported to date. However, the highly conserved molecular architecture and sequence motifs that characterize the class A GPCR family make it possibly to reliably predict the topological positions of the identified mutations from structural-informed sequence alignments. Using this approach, we provide a 3D context for the many variants occurring in ORs facilitating the functional interpretation of the changes attending to their structural location, biochemical associated data and substitution score weightings. This approach is exemplified through the identification of several OR natural variants located at conserved topological sites (e.g. BW 2.50, 3.50, 7.50, 45.50 at ECL2), either involved in the structural stability, or in the functional mechanism of the receptors, and which might induce changes in the odorant sensitivity. We believe the integration of high-throughput sequencing data with structural information is crucial for the interpretation of the complex genotype-phenotype associations occurring not only in human olfaction, but also in any other biological process. These would require in many cases the development of automatic interfaces to facilitate the management and organization of large quantities of data. Hence, we develop an interactive computational application that integrates both, the genomic and structural knowledge with analytical graphical tools, for the study of the OR mutational landscape. The hORMdb, available at (http://lmc.uab.cat/hORMdb) allows the comparison, topological localization and evaluation of natural variations occurred in human ORs, and represents to our knowledge, one of the largest collections of variation data of human sensory proteins annotated at structural level. We envisage that the utility of this information will increase as the amount of available pharmacological data for these receptors grow. This effort, together with ongoing research in the study of genetic changes in other sensory receptors [63] could shape an emerging *sensegenomics* field of knowledge, which should be considered by food and cosmetic consumer product manufacturers in benefit of the general population.

## AVAILABILITY

The human Olfactory Receptor Mutation database is an open source initiative available in the GitHub repository (https://github.com/lmc-uab/hORMdb).

## SUPPLEMENTARY DATA

Supplementary Data are available at BioRxiv online.

