## Supplemental Figures and Tables for "The mutational landscape of human olfactory G protein-coupled receptors"

### Supplementary Figures and Tables:

- **Figure S1:** Multiple sequence alignment (MSA) of human ORs.
- **Figure S2:** Structure-based sequence alignment used for topological annotation.
- **Figure S3:** Functional core (FC) of topological positions in class A GPCRs.
- **Figure S4:** Binding site (BS) topological positions in class A GPCRs.
- **Figure S5:** The GPCRtm substitution matrix.
- **Figure S6:** Chromosomal distribution of natural variants within OR families.
- **Figure S7:** Topological distribution of natural variants within OR families.
- **Figure S8:** Human OR mutations with potential functional effects.

- **Table S1:** Nucleotide sequencing data sources used in the study.
- **Table S2:** Number of mutations in human ORs collected in the study.
- **Table S3:** Conserved topological sites with functional implication in the GPCR activity.
- **Table S4:** Non-olfactory class A GPCRs used in topological annotation.
- **Table S5:** The human OR mutation database table.

**Supplementary Figure S1: Multiple sequence alignment (MSA) of human ORs.** The resulting MSA of the 374 human OR UniProt sequences used in the topological annotation of protein-coding mutations can be found at http://lmc.uab.cat/hORMdb. Receptor sequences were aligned with ClustalW (v2.1) using a customized GPCR substitution score matrix [1]. The resulted MSA was manually adjusted to fulfil the structural information derived from non-olfactory class A GPCRs (Supplementary Figure S2). Topological regions (N- and C-terminal sequences, transmembrane α-helices TM 1 to 7, extracellular ECL 1 to 3 and cytoplasmic loops ICL 1 to 3), as well as Ballesteros-Weinstein (BW), functional core (FC) and ligand binding site (BS) topological positions are indicated on top of the alignment.

**Supplementary Figure S2:**

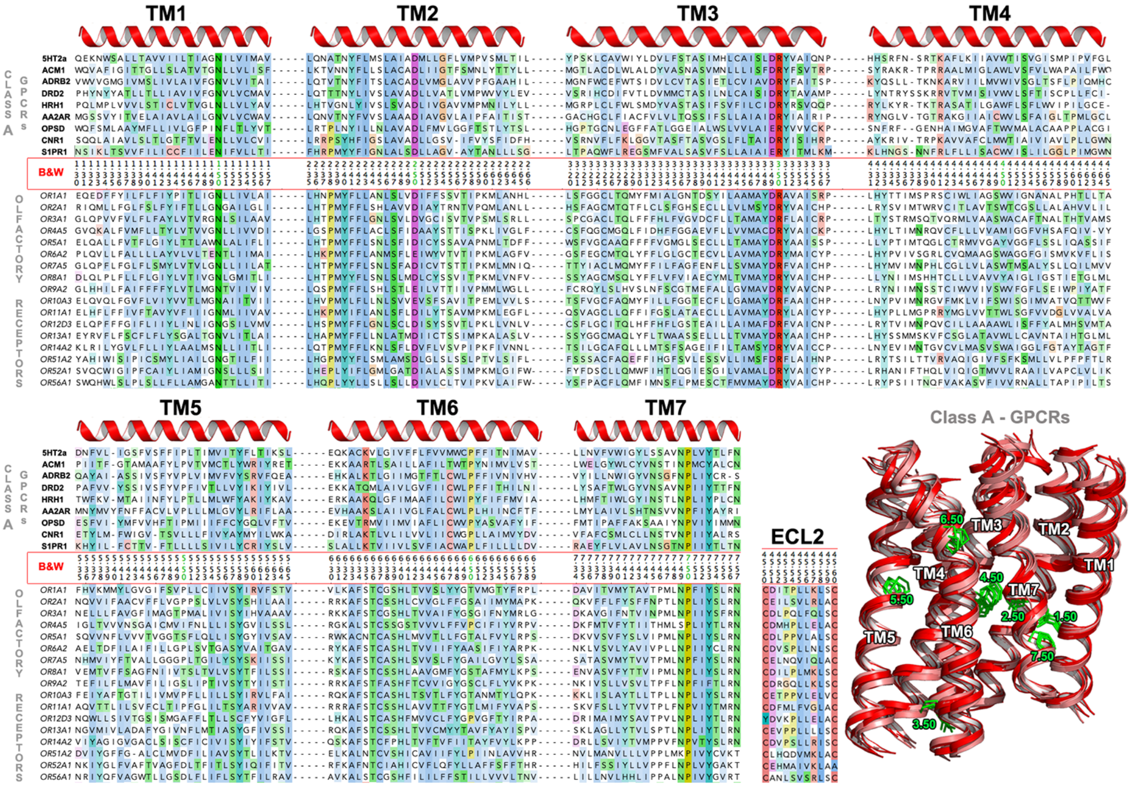


**Supplementary Figure S2: Structure-based sequence alignment used for topological annotation.** Sequence alignment of the TM regions of representative non-olfactory class A GPCRs with known 3D-structures and one member of each of the 17 OR families analyzed (the complete lists of receptors are available in Supplementary Tables 2 and 4). Non-olfactory receptors in the figure correspond to 5-hydroxytryptamine 2A (5HT2a, PDBid: 6A94), acetylcholine muscarinic (ACM1, PDBid: 6OIJ), 2-beta adrenergic (ADRB2, PDBid: 5JQH), dopamine D2 (DRD2, PDBid: 6CM4), histamine H1 (HRH1, PDBid: 3RZE), adenosine A2A (AA2AR, PDBid: 3VG9), rhodopsin (OPSD, PDBid: 1GZM), cannabinoid (CNR1, PDBid: 5TGZ) and Sphingosine 1-phosphate receptor (S1PR1, PDBid: 3V2Y). On the lower right is shown the structural superimposition of their TM regions with the most conserved BW positions (.50) at each helix highlighted. Ribbon diagrams on top of the alignment indicate the boundaries of the TM regions according to the structural superposition. The red frame in the alignment indicates BW positions [2]. An adaptation of this numbering system was applied for a conserved stretch of 10 residues at the ECL2 (indicated by 45 as first number attending to its location between the TMs 4 and 5) [3].

**Supplementary Figure S3:**

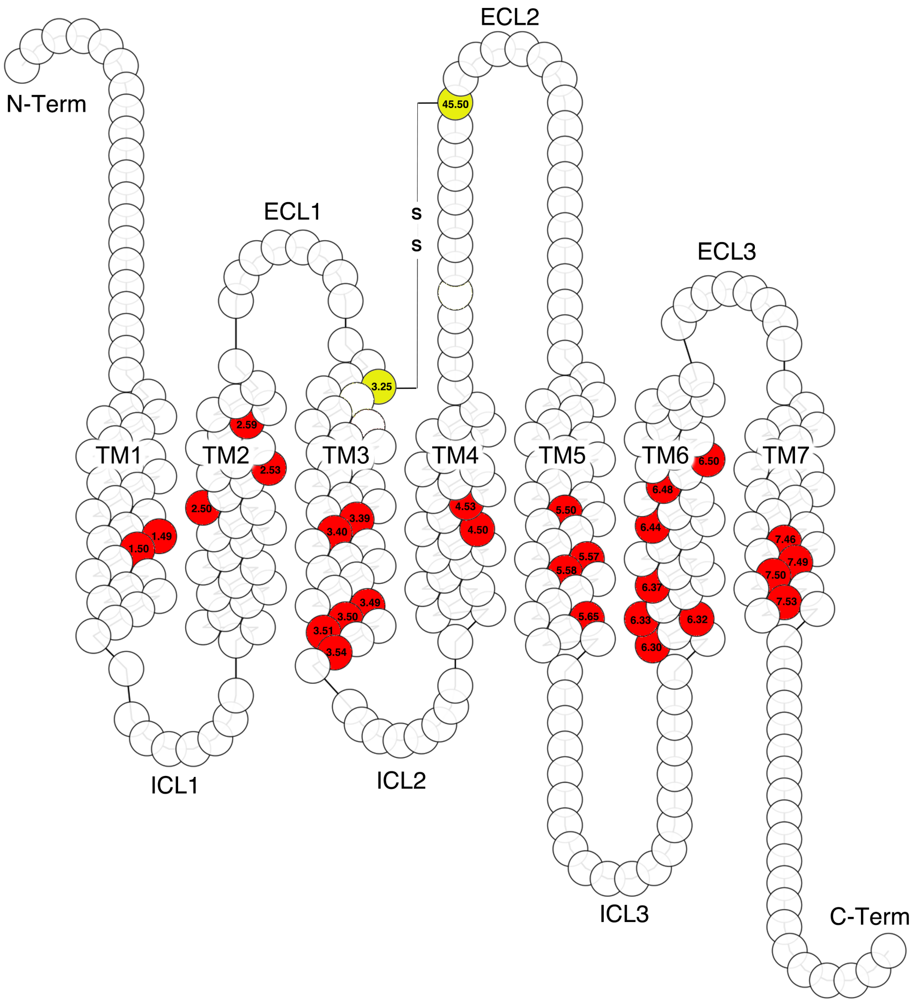


**Supplementary Figure S3: Functional core (FC) topological positions in class A GPCRs.** Snake plot representation of a generic class A GPCR with topological regions labeled. Color filled circles with BW notation indicate positions likely involved in the receptor activation or G-protein interaction (red) and conserved cysteines forming part of disulfide bridges (in yellow). Residue conservation and reference information to each position is available in the Supplementary Table S3.

**Supplementary Figure S4:**

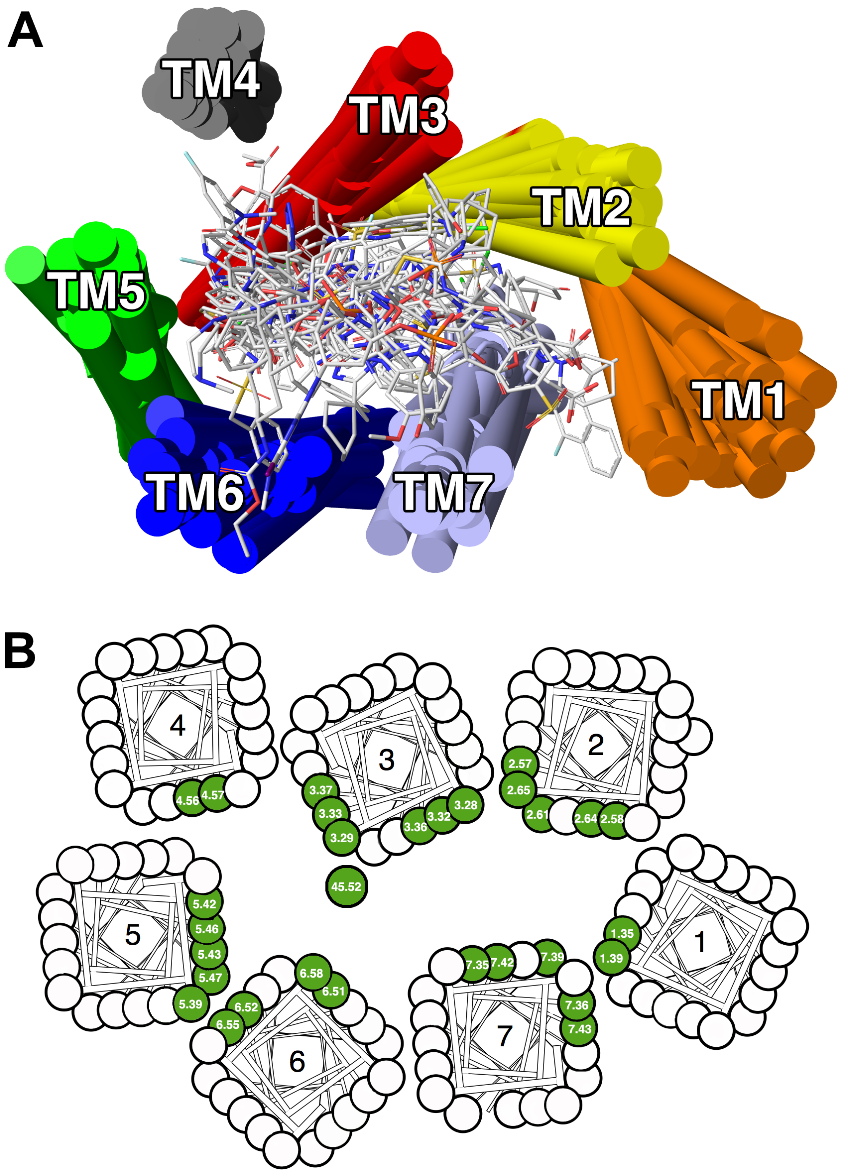


**Supplementary Figure S4: Binding site topological positions in class A GPCRs.** A. Structural superimposition of 39 class A GPCRs with known three-dimensional structures used for a general binding site (BS) definition (the complete list of the receptors is available in the Supplementary Table S4). The molecular coordinates of the TM regions of each receptor with their corresponding ligands are represented in color tubes and sticks, respectively. B. Snake plot representation of the extracellular view of a generic GPCR with the TM regions indicated by numbers. Color filled circles with BW notation indicate the positions within a distance ≤ 4.0 Å of ligands in the crystallographic structures displayed in A.

**Supplementary Figure S5:**

| | A | C | D | E | F | G | H | I | K | L | M | N | P | Q | R | S | T | V | W | Y |
|---|---|---|---|---|---|---|---|---|---|---|---|---|---|---|---|---|---|---|---|---|
| **A** | 2 | | | | | | | | | | | | | | | | | | | |
| **C** | 0 | 5 | | | | | | | | | | | | | | | | | | |
| **D** | -3 | -3 | 9 | | | | | | | | | | | | | | | | | |
| **E** | -1 | -3 | 5 | 6 | | | | | | | | | | | | | | | | |
| **F** | -1 | 0 | -2 | -2 | 2 | | | | | | | | | | | | | | | |
| **G** | 1 | 0 | -1 | 1 | 0 | 4 | | | | | | | | | | | | | | |
| **H** | -1 | -1 | 1 | 1 | -1 | -1 | 5 | | | | | | | | | | | | | |
| **I** | -1 | -1 | -4 | -2 | 0 | -1 | -1 | 2 | | | | | | | | | | | | |
| **K** | -1 | -2 | 0 | 3 | -1 | -1 | 2 | -2 | 6 | | | | | | | | | | | |
| **L** | -1 | -1 | -4 | -2 | 0 | -1 | -1 | 1 | -2 | 2 | | | | | | | | | | |
| **M** | -1 | 0 | -3 | -1 | 0 | 0 | 0 | 1 | -1 | 1 | 3 | | | | | | | | | |
| **N** | -2 | -3 | 2 | 0 | -2 | -2 | 2 | -3 | 0 | -3 | -2 | 8 | | | | | | | | |
| **P** | -1 | -3 | 0 | 0 | -3 | -2 | 0 | -3 | -3 | -3 | -3 | -2 | 8 | | | | | | | |
| **Q** | -1 | -2 | 1 | 3 | -1 | -1 | 3 | -2 | 3 | -1 | -1 | 1 | 0 | 5 | | | | | | |
| **R** | -1 | -1 | -1 | 2 | -2 | -1 | 2 | -3 | 4 | -2 | -2 | 0 | -2 | 3 | 7 | | | | | |
| **S** | 1 | 0 | -2 | 1 | -1 | 1 | 0 | -1 | 0 | -1 | -1 | 0 | 0 | 0 | -1 | 2 | | | | |
| **T** | 1 | 0 | -2 | -1 | -1 | 1 | 0 | 0 | -1 | 0 | 0 | -1 | -1 | -1 | -1 | 1 | 2 | | | |
| **V** | 0 | 0 | -3 | -2 | 0 | -1 | -1 | 1 | -2 | 0 | 0 | -3 | -3 | -1 | -2 | -1 | 0 | 1 | | |
| **W** | -2 | -1 | -3 | -1 | 0 | -1 | 0 | -2 | -1 | -1 | -1 | -2 | -4 | 1 | 0 | -2 | -2 | -1 | 8 | |
| **Y** | -2 | -1 | -2 | -1 | 1 | -1 | 1 | -2 | -1 | -1 | -1 | -1 | -4 | 0 | -2 | -2 | -2 | -1 | 2 | 6 |

**Supplementary Figure S5: The GPCRtm amino acid substitution scores.** Values in the matrix correspond to statistical amino acid substitution scores calculated for the 20 aminoacids (one-letter code) in a MSA of more than one thousand class A GPCR sequences including human ORs [1].

**Supplementary Figure S6:**

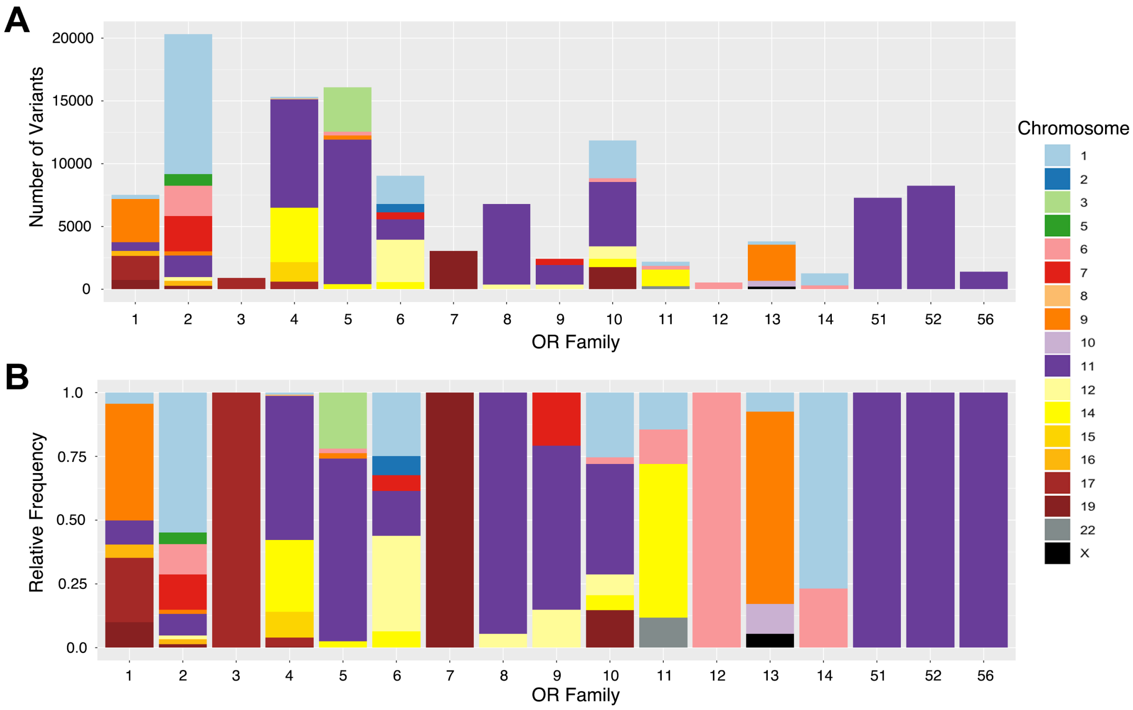


**Supplementary Figure S6: Chromosomal distribution of natural variants within OR families.** A. Total number of collected variants (y-axis) at each to the 17 OR families analyzed (x-axis). Bars are colored according to the chromosomal location of the natural variants (color legend on the right). B. Relative frequencies of the chromosomal distribution of the mutations at each OR family.

**Supplementary Figure S7:**

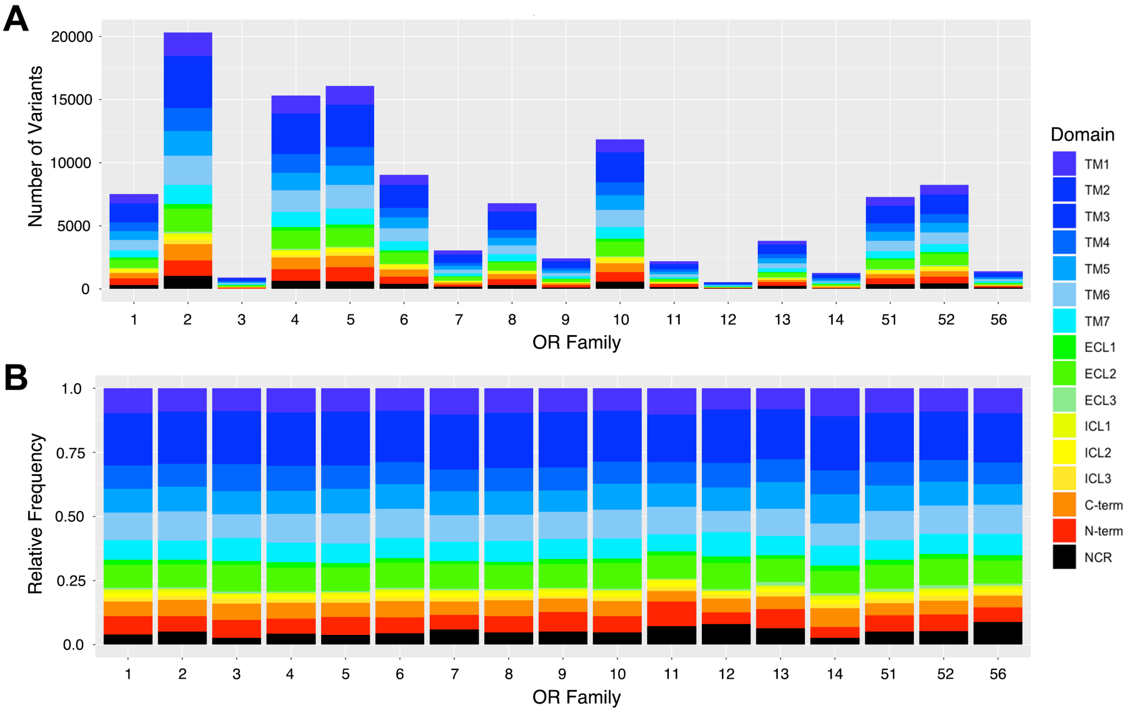


**Supplementary Figure S7: Topological distribution of natural variants within OR families.** A. Total number of collected variants (y-axis) at each to the 17 OR families analyzed (x-axis) color-coded by their GPCR domain location (color legend on the right). B. Relative frequencies of the topological domain distribution of the natural variants at each OR family.

**Supplementary Figure S8:**

| rsID | Mutation | Role | Gene | Allele frequencies (%) AFR | ASH | EF | ENF | LAT | EA | SA |
|---|---|---|---|---|---|---|---|---|---|---|
| rs12800642 | V1.35L | BS | OR4C16 | 13 | 31 | 20 | 22 | 20 | 14 | 21 |
| rs73146195 | G1.49E | FC | OR2T4 | 25 | 2 | 8 | 4 | 4 | 9 | 5 |
| rs4501959 | D2.50N | FC | OR52L1 | 4 | 20 | 17 | 20 | 39 | 33 | 11 |
| rs4388990 | Y2.57C | BS | OR6C74 | 18 | 34 | 29 | 31 | 17 | 12 | 34 |
| rs12365487 | T2.57N | BS | OR52N1 | 5 | 16 | 10 | 11 | 17 | 25 | 19 |
| rs62006708 | A2.58P | BS | OR4N4 | 4 | 24 | 21 | 22 | 10 | 18 | 23 |
| rs7305779 | E2.58A | BS | OR9K2 | 40 | 34 | 29 | 32 | 18 | 11 | 35 |
| rs6658227 | P2.59L | FC | OR2L3 | 27 | 8 | 14 | 14 | 7 | 27 | 9 |
| rs6664332 | S2.64N | BS | OR2AK2 | 28 | 26 | 34 | 34 | 36 | 46 | 45 |
| rs61818749 | C3.25Y | FC | OR10T2 | 2 | 5 | 10 | 10 | 6 | 13 | 18 |
| rs76383258 | Q3.28L | FC | OR5D14 | 14 | 12 | 6 | 11 | 13 | 24 | 30 |
| rs12118628 | M3.29I | BS | OR10J1 | 4 | 12 | 11 | 12 | 27 | 23 | 13 |
| rs11036912 | I3.32T | BS | OR51B5 | 27 | 40 | 49 | 46 | 41 | 17 | 40 |
| rs139616012 | H3.32Y | BS | OR2T34 | 10 | 2 | 1 | 2 | 5 | 9 | 3 |
| rs28540568 | V3.33L | BS | OR2T4 | 18 | 12 | 19 | 15 | 30 | 44 | 33 |
| rs117277709 | G3.36V | BS | OR5L2 | 8 | 1 | 3 | 3 | 2 | 4 | 1 |
| rs61750896 | S3.40L | FC | OR52L1 | 15 | 9 | 7 | 9 | 6 | 11 | 14 |
| rs10836954 | S3.40L | FC | OR51G1 | 1 | 7 | 9 | 6 | 2 | 24 | 5 |
| rs2073153 | L3.50R | FC | OR12D2 | 20 | 49 | 42 | 44 | 39 | 33 | 28 |
| rs77989044 | C3.50R | FC | OR51B6 | 23 | 10 | 7 | 5 | 11 | 5 | 10 |
| rs74396937 | H3.50R | FC | OR4C46 | 16 | 11 | 6 | 10 | 13 | 23 | 28 |
| rs12224086 | R3.50L | FC | OR5AS1 | 5 | 9 | 19 | 12 | 11 | 16 | 13 |
| rs10176036 | R3.50C | FC | OR6B2 | 11 | 27 | 42 | 32 | 29 | 40 | 17 |
| rs3751484 | R3.50H | FC | OR6J1 | 32 | 10 | 12 | 18 | 12 | 17 | 14 |
| rs11230983 | R3.50H | FC | OR5D13 | 5 | 9 | 19 | 12 | 11 | 16 | 12 |
| rs2512219 | R3.50H | FC | OR8D2 | 43 | 23 | 15 | 15 | 18 | 18 | 32 |
| rs16930982 | R3.50H | FC | OR51I1 | 1 | 5 | 1 | 2 | 11 | 38 | 13 |
| rs2072164 | R3.50C | FC | OR2F1 | 8 | 6 | 13 | 6 | 24 | 17 | 11 |
| rs11227321 | Y3.51H | FC | OR8J3 | 5 | 9 | 19 | 12 | 11 | 16 | 13 |
| rs10769224 | C3.51Y | FC | OR52N1 | 44 | 30 | 31 | 30 | 27 | 22 | 27 |
| rs2073152 | S3.51C | FC | OR12D2 | 12 | 32 | 28 | 29 | 15 | 23 | 12 |
| rs10839659 | W4.50S | FC | OR2D3 | 9 | 35 | 37 | 38 | 30 | 36 | 32 |
| rs73406604 | Y4.50C | FC | OR5P2 | 34 | 23 | 28 | 28 | 29 | 13 | 15 |
| rs28556931 | M4.50L | FC | OR13G1 | 30 | 32 | 31 | 36 | 22 | 39 | 36 |
| rs138825902 | D4.56N | BS | OR2T27 | 18 | 5 | 10 | 8 | 8 | 24 | 3 |
| rs199920874 | D4.56N | BS | OR2T7 | 11 | 5 | 12 | 7 | 7 | 15 | 12 |
| rs10187574 | C45.50R | FC | OR6B2 | 17 | 28 | 44 | 34 | 31 | 43 | 18 |
| rs58664826 | C45.50Y | FC | OR52J3 | 4 | 12 | 10 | 13 | 31 | 11 | 7 |
| rs4057749 | C45.50R | FC | OR8B4 | 41 | 27 | 20 | 22 | 14 | 22 | 26 |
| rs34337292 | C45.50R | FC | OR9Q2 | 28 | 16 | 23 | 27 | 32 | 7 | 17 |
| rs9901356 | L45.52V | BS | OR3A2 | 29 | 16 | 12 | 12 | 23 | 6 | 28 |
| rs4584426 | T45.52A | BS | OR2T8 | 20 | 21 | 24 | 30 | 26 | 20 | 13 |
| rs142669752 | M5.39R | BS | OR2T33 | 3 | 21 | 18 | 21 | 17 | 6 | 12 |
| rs7970885 | V5.39M | BS | OR10P1 | 15 | 44 | 41 | 46 | 39 | 29 | 31 |
| rs12221615 | L5.39V | BS | OR5T2 | 6 | 9 | 19 | 12 | 11 | 15 | 13 |
| rs79101657 | S5.42C | BS | OR4M2 | 2 | 2 | 2 | 2 | 3 | 13 | 2 |
| rs2466584 | N5.42S | BS | OR10D3 | 11 | 22 | 14 | 14 | 10 | 15 | 29 |
| rs61738869 | C5.43G | BS | OR10H5 | 5 | 14 | 5 | 10 | 6 | 8 | 6 |
| rs9822460 | I5.46V | BS | OR5K4 | 26 | 13 | 25 | 21 | 18 | 15 | 26 |
| rs513873 | V5.47A | BS | OR9G4 | 24 | 37 | 21 | 32 | 16 | 8 | 19 |
| rs12363178 | W5.47G | BS | OR52N4 | 15 | 20 | 14 | 17 | 19 | 24 | 23 |
| rs10466659 | V5.50A | FC | OR5B2 | 23 | 22 | 25 | 22 | 17 | 14 | 25 |
| rs16979912 | S5.50P | FC | OR7C1 | 42 | 11 | 18 | 19 | 25 | 3 | 20 |
| rs76615008 | C5.57S | FC | OR5P2 | 34 | 23 | 28 | 28 | 29 | 13 | 15 |
| rs11880955 | I5.65L | FC | OR7A10 | 23 | 15 | 15 | 17 | 12 | 7 | 11 |
| rs16839214 | I5.65V | FC | OR5H2 | 15 | 22 | 9 | 12 | 25 | 26 | 13 |
| rs17350764 | V5.65I | FC | OR52J3 | 25 | 22 | 14 | 19 | 35 | 12 | 26 |
| rs35393723 | R6.30W | FC | OR10J5 | 4 | 12 | 25 | 19 | 15 | 1 | 16 |
| rs11265165 | Q6.30R | FC | OR10J3 | 16 | 12 | 11 | 8 | 14 | 46 | 13 |
| rs61833441 | R6.30H | FC | OR2T34 | 22 | 11 | 15 | 11 | 23 | 29 | 22 |
| rs2570571 | E6.30Q | FC | OR51A2 | 14 | 29 | 35 | 29 | 17 | 14 | 25 |
| rs67863827 | R6.30H | FC | OR5D14 | 13 | 7 | 6 | 9 | 4 | 8 | 6 |
| rs10838881 | K6.32R | FC | OR4C5 | 28 | 13 | 24 | 16 | 15 | 18 | 17 |
| rs4936880 | R6.32G | FC | OR10G4 | 48 | 49 | 43 | 46 | 48 | 35 | 38 |
| rs78350717 | A6.33V | FC | OR11H1 | 10 | 32 | 32 | 36 | 18 | 25 | 19 |
| rs7101919 | I6.44T | FC | OR52D1 | 9 | 25 | 37 | 33 | 26 | 4 | 19 |
| rs7106330 | V6.52L | BS | OR51B6 | 5 | 8 | 14 | 16 | 19 | 8 | 15 |
| rs74053516 | L6.52I | BS | OR52B6 | 3 | 6 | 3 | 5 | 2 | 1 | 1 |
| rs11823842 | F6.52L | BS | OR52E4 | 33 | 33 | 39 | 43 | 36 | 36 | 34 |
| rs199863869 | M7.35V | BS | OR2T34 | 30 | 10 | 14 | 12 | 10 | 11 | 12 |
| rs7934354 | M7.36T | BS | OR52H1 | 14 | 4 | 6 | 2 | 10 | 36 | 12 |
| rs2512166 | T7.42A | BS | OR8G5 | 3 | 14 | 13 | 15 | 14 | 11 | 10 |
| rs7130086 | T7.42K | BS | OR4B1 | 15 | 23 | 14 | 18 | 15 | 36 | 43 |
| rs11880184 | T7.42A | BS | OR10H4 | 43 | 27 | 26 | 25 | 14 | 9 | 17 |
| rs2512166 | T7.42A | BS | OR8G1 | 3 | 14 | 13 | 15 | 14 | 11 | 10 |
| rs4936845 | T7.42K | BS | OR6M1 | 11 | 32 | 31 | 29 | 32 | 32 | 27 |
| rs1681596 | V7.43L | BS | OR6J1 | 33 | 13 | 15 | 22 | 14 | 17 | 15 |
| rs11036815 | P7.46S | FC | OR51B2 | 29 | 33 | 27 | 28 | 27 | 21 | 25 |
| rs769427 | P7.50S | FC | OR1A1 | 36 | 46 | 46 | 49 | 28 | 14 | 44 |
| rs12790505 | S7.50P | FC | OR5L1 | 16 | 7 | 6 | 9 | 4 | 8 | 6 |

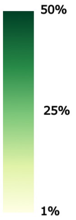


**Supplementary Figure S8: Human OR mutations with potential functional effects.** Eighty natural variants selected from the study with allele frequencies > 1% and belonging to FC and BS topological regions (see methods). For each natural variant a reference rsID number, BW position, type of substitution, functional region and gene ID are provided. Colored boxes indicate the allele frequencies expressed as percentage (color legend on the right) in seven sub-continental populations: AFR (African),  ASH - Ashkenazi Jewish, European Finnish (EF), East Asian (EA), (), Latino (LAT), South Asian (SA) and European non-Finnish (ENF).

### Supplementary Table S1

| Population | Abbr. | Genomes | Exomes | Total |
|---|---|---|---|---|
| African/African-American | AFR | 4.359 | 8,128 | 12,487 |
| Admixed American (Latino) | LAT | 419 | 16,791 | 17,210 |
| Ashkenazi Jewish | ASH | 145 | 5,040 | 5,185 |
| East Asian | EA | 780 | 9,197 | 9,977 |
| European Finnish | EF | 1,738 | 10,824 | 12,562 |
| European Non-Finnish | ENF | 7,718 | 56,885 | 64,603 |
| South Asian | SA | | 15,308 | 15,308 |
| Other (population not assigned) | OTH | 544 | 3,070 | 3,614 |
| **Total** | | **15,708** | **125,748** | **141,456** |

**Supplementary Table S1: Nucleotide sequencing data sources used in the study.** Human OR mutation data was taken from the genome aggregation database (gnomAD), comprising a total of 16 million single nucleotide variation (SNVs) and 1.2 million indels from 125.748 exomes, and 229 million SNVs and 33 million indels from 15.708 genomes (https://macarthurlab.org/2017/02/27/the-genome-aggregation-database-gnomad/). Samples were subdivided in six geographic ancestries according to a random forest classifier using principal component analysis (PCA), plus a eighth group named “Other” (OTH) that include individuals that do not unambiguously cluster within any of the foregoing populations [4, 5].

### Supplementary Table S2

| OR Family 1 | | OR Family 2 | | OR Family 3 | | OR Family 4 | | OR Family 5 | |
|---|---|---|---|---|---|---|---|---|---|
| OR Name | № Mut | OR Name | № Mut | OR Name | № Mut | OR Name | № Mut | OR Name | № Mut |
| OR1A1 | 292 | OR2A1 | 90 | OR3A1 | 328 | OR4A15 | 566 | OR5A1 | 350 |
| OR1A2 | 241 | OR2A12 | 304 | OR3A2 | 329 | OR4A16 | 552 | OR5A2 | 312 |
| OR1B1 | 311 | OR2A14 | 333 | OR3A3 | 253 | OR4A47 | 453 | OR5AC2 | 341 |
| OR1C1 | 334 | OR2A2 | 312 | | | OR4A5 | 624 | OR5AK2 | 371 |
| OR1D2 | 278 | OR2A25 | 318 | | | OR4B1 | 391 | OR5AN1 | 324 |
| OR1D5 | 244 | OR2A4 | 197 | | | OR4C11 | 355 | OR5AP2 | 293 |
| OR1E1 | 257 | OR2A5 | 323 | | | OR4C12 | 224 | OR5AR1 | 336 |
| OR1E2 | 296 | OR2A7 | 158 | | | OR4C13 | 323 | OR5AS1 | 338 |
| OR1F1 | 389 | OR2AE1 | 327 | | | OR4C15 | 33 | OR5AU1 | 405 |
| OR1G1 | 295 | OR2AG1 | 337 | | | OR4C16 | 514 | OR5B12 | 313 |
| OR1I1 | 372 | OR2AG2 | 373 | | | OR4C3 | 29 | OR5B17 | 301 |
| OR1J1 | 325 | OR2AJ1 | 207 | | | OR4C46 | 557 | OR5B2 | 332 |
| OR1J2 | 295 | OR2AK2 | 277 | | | OR4C5 | 229 | OR5B21 | 293 |
| OR1J4 | 259 | OR2AP1 | 297 | | | OR4C6 | 425 | OR5B3 | 314 |
| OR1K1 | 346 | OR2AT4 | 294 | | | OR4D1 | 267 | OR5C1 | 352 |
| OR1L1 | 51 | OR2B11 | 334 | | | OR4D10 | 316 | OR5D13 | 339 |
| OR1L3 | 343 | OR2B2 | 267 | | | OR4D11 | 307 | OR5D14 | 344 |
| OR1L4 | 280 | OR2B3 | 266 | | | OR4D2 | 279 | OR5D16 | 350 |
| OR1L6 | 47 | OR2B6 | 238 | | | OR4D5 | 324 | OR5D18 | 294 |
| OR1L8 | 272 | OR2C1 | 386 | | | OR4D6 | 315 | OR5F1 | 357 |
| OR1M1 | 379 | OR2C3 | 342 | | | OR4D9 | 333 | OR5H1 | 355 |
| OR1N1 | 291 | OR2D2 | 369 | | | OR4E2 | 306 | OR5H14 | 404 |
| OR1N2 | 320 | OR2D3 | 342 | | | OR4F15 | 308 | OR5H15 | 375 |
| OR1Q1 | 295 | OR2F1 | 310 | | | OR4F17 | 58 | OR5H2 | 292 |
| OR1S1 | 363 | OR2F2 | 340 | | | OR4F21 | 56 | OR5H6 | 430 |
| OR1S2 | 345 | OR2G2 | 315 | | | OR4F4 | 179 | OR5I1 | 320 |
| | | OR2G3 | 275 | | | OR4F5 | 132 | OR5J2 | 328 |
| | | OR2G6 | 412 | | | OR4F6 | 320 | OR5K1 | 352 |
| | | OR2H1 | 308 | | | OR4K1 | 392 | OR5K2 | 333 |
| | | OR2H2 | 305 | | | OR4K13 | 309 | OR5K3 | 331 |
| | | OR2J2 | 298 | | | OR4K14 | 283 | OR5K4 | 316 |
| | | OR2J3 | 260 | | | OR4K15 | 404 | OR5L1 | 371 |
| | | OR2K2 | 29 | | | OR4K17 | 21 | OR5L2 | 351 |
| | | OR2L13 | 325 | | | OR4K2 | 382 | OR5M1 | 345 |
| | | OR2L2 | 366 | | | OR4K5 | 365 | OR5M10 | 381 |
| | | OR2L3 | 315 | | | OR4L1 | 345 | OR5M11 | 334 |
| | | OR2L5 | 342 | | | OR4M1 | 413 | OR5M3 | 349 |
| | | OR2L8 | 312 | | | OR4M2 | 362 | OR5M8 | 372 |
| | | OR2M2 | 349 | | | OR4N2 | 378 | OR5M9 | 347 |
| | | OR2M3 | 345 | | | OR4N4 | 378 | OR5P2 | 400 |
| | | OR2M4 | 275 | | | OR4N5 | 322 | OR5P3 | 315 |
| | | OR2M5 | 384 | | | OR4P4 | 275 | OR5R1 | 319 |
| | | OR2M7 | 315 | | | OR4Q3 | 412 | OR5T1 | 369 |
| | | OR2S2 | 323 | | | OR4S1 | 379 | OR5T2 | 416 |
| | | OR2T1 | 386 | | | OR4S2 | 265 | OR5T3 | 315 |
| | | OR2T10 | 284 | | | OR4X1 | 394 | OR5V1 | 282 |
| | | OR2T11 | 381 | | | OR4X2 | 470 | OR5W2 | 315 |
| | | OR2T12 | 382 | | | | | | |
| | | OR2T2 | 400 | | | | | | |
| | | OR2T27 | 425 | | | | | | |
| | | OR2T29 | 106 | | | | | | |
| | | OR2T3 | 351 | | | | | | |
| | | OR2T33 | 390 | | | | | | |
| | | OR2T34 | 350 | | | | | | |
| | | OR2T35 | 257 | | | | | | |
| | | OR2T4 | 367 | | | | | | |
| | | OR2T5 | 101 | | | | | | |
| | | OR2T6 | 322 | | | | | | |
| | | OR2T7 | 451 | | | | | | |
| | | OR2T8 | 309 | | | | | | |
| | | OR2V1 | 286 | | | | | | |
| | | OR2V2 | 297 | | | | | | |
| | | OR2W1 | 263 | | | | | | |
| | | OR2W3 | 382 | | | | | | |
| | | OR2Y1 | 352 | | | | | | |
| | | OR2Z1 | 276 | | | | | | |

| OR Family 6 | | OR Family 7 | | OR Family 8 | | OR Family 9 | | OR Family 10 | |
|---|---|---|---|---|---|---|---|---|---|
| OR Name | № Mut | OR Name | № Mut | OR Name | № Mut | OR Name | № Mut | OR Name | № Mut |
| OR6A2 | 348 | OR7A10 | 306 | OR8A1 | 315 | OR9A2 | 266 | OR10A2 | 309 |
| OR6B1 | 293 | OR7A17 | 345 | OR8B12 | 284 | OR9A4 | 239 | OR10A3 | 326 |
| OR6B2 | 349 | OR7A5 | 286 | OR8B2 | 295 | OR9G1 | 246 | OR10A4 | 332 |
| OR6B3 | 322 | OR7C1 | 284 | OR8B3 | 297 | OR9G4 | 353 | OR10A5 | 338 |
| OR6C1 | 310 | OR7C2 | 278 | OR8B4 | 261 | OR9I1 | 291 | OR10A6 | 382 |
| OR6C2 | 315 | OR7D2 | 299 | OR8B8 | 314 | OR9K2 | 359 | OR10A7 | 299 |
| OR6C3 | 352 | OR7D4 | 339 | OR8D1 | 291 | OR9Q1 | 307 | OR10AD1 | 337 |
| OR6C4 | 356 | OR7E24 | 291 | OR8D2 | 261 | OR9Q2 | 358 | OR10AG1 | 316 |
| OR6C6 | 300 | OR7G1 | 319 | OR8D4 | 296 | | | OR10C1 | 306 |
| OR6C65 | 280 | OR7G2 | 12 | OR8G1 | 339 | | | OR10D3 | 239 |
| OR6C68 | 336 | OR7G3 | 284 | OR8G5 | 332 | | | OR10G2 | 381 |
| OR6C70 | 309 | | | OR8H1 | 344 | | | OR10G3 | 295 |
| OR6C74 | 279 | | | OR8H2 | 328 | | | OR10G4 | 324 |
| OR6C75 | 257 | | | OR8H3 | 343 | | | OR10G6 | 100 |
| OR6C76 | 285 | | | OR8I2 | 345 | | | OR10G7 | 362 |
| OR6F1 | 296 | | | OR8J1 | 307 | | | OR10G8 | 352 |
| OR6J1 | 240 | | | OR8J3 | 315 | | | OR10G9 | 372 |
| OR6K2 | 372 | | | OR8K1 | 323 | | | OR10H1 | 390 |
| OR6K3 | 31 | | | OR8K3 | 321 | | | OR10H2 | 361 |
| OR6K6 | 346 | | | OR8K5 | 295 | | | OR10H3 | 295 |
| OR6M1 | 312 | | | OR8S1 | 374 | | | OR10H4 | 304 |
| OR6N1 | 319 | | | OR8U1 | 208 | | | OR10H5 | 404 |
| OR6N2 | 274 | | | | | | | OR10J1 | 407 |
| OR6P1 | 280 | | | | | | | OR10J3 | 331 |
| OR6Q1 | 301 | | | | | | | OR10J5 | 291 |
| OR6S1 | 344 | | | | | | | OR10K1 | 337 |
| OR6T1 | 358 | | | | | | | OR10K2 | 303 |
| OR6V1 | 267 | | | | | | | OR10P1 | 343 |
| OR6X1 | 283 | | | | | | | OR10Q1 | 389 |
| OR6Y1 | 332 | | | | | | | OR10R2 | 327 |
| | | | | | | | | OR10S1 | 363 |
| | | | | | | | | OR10T2 | 319 |
| | | | | | | | | OR10V1 | 305 |
| | | | | | | | | OR10W1 | 332 |
| | | | | | | | | OR10X1 | 348 |
| | | | | | | | | OR10Z1 | 339 |

| OR Family 11 | | OR Family 12 | | OR Family 13 | | OR Family 14 | | OR Family 51 | | OR Family 52 | | OR Family 56 | |
|---|---|---|---|---|---|---|---|---|---|---|---|---|---|
| OR Name | № Mut | OR Name | № Mut | OR Name | № Mut | OR Name | № Mut | OR Name | № Mut | OR Name | № Mut | OR Name | № Mut |
| OR11A1 | 296 | OR12D2 | 281 | OR13A1 | 450 | OR14A16 | 256 | OR51A2 | 216 | OR52A1 | 310 | OR56A1 | 329 |
| OR11G2 | 314 | OR12D3 | 259 | OR13C2 | 359 | OR14A2 | 183 | OR51A4 | 295 | OR52A5 | 306 | OR56A3 | 300 |
| OR11H1 | 256 | | | OR13C3 | 336 | OR14C36 | 288 | OR51A7 | 282 | OR52B2 | 335 | OR56A4 | 47 |
| OR11H12 | 331 | | | OR13C4 | 272 | OR14J1 | 296 | OR51B2 | 328 | OR52B4 | 387 | OR56B1 | 363 |
| OR11H4 | 354 | | | OR13C5 | 363 | OR14K1 | 255 | OR51B4 | 327 | OR52B6 | 297 | OR56B4 | 350 |
| OR11H6 | 320 | | | OR13C8 | 274 | | | OR51B5 | 412 | OR52D1 | 358 | | |
| OR11L1 | 316 | | | OR13C9 | 282 | | | OR51B6 | 357 | OR52E2 | 308 | | |
| | | | | OR13D1 | 324 | | | OR51D1 | 381 | OR52E4 | 329 | | |
| | | | | OR13F1 | 325 | | | OR51E1 | 39 | OR52E6 | 361 | | |
| | | | | OR13G1 | 286 | | | OR51E2 | 392 | OR52E8 | 372 | | |
| | | | | OR13H1 | 210 | | | OR51F1 | 311 | OR52H1 | 312 | | |
| | | | | OR13J1 | 350 | | | OR51F2 | 354 | OR52I1 | 348 | | |
| | | | | | | | | OR51G1 | 375 | OR52I2 | 345 | | |
| | | | | | | | | OR51G2 | 364 | OR52J3 | 337 | | |
| | | | | | | | | OR51H1 | 284 | OR52K1 | 412 | | |
| | | | | | | | | OR51I1 | 350 | OR52K2 | 408 | | |
| | | | | | | | | OR51I2 | 384 | OR52L1 | 338 | | |
| | | | | | | | | OR51L1 | 279 | OR52M1 | 485 | | |
| | | | | | | | | OR51M1 | 394 | OR52N1 | 290 | | |
| | | | | | | | | OR51Q1 | 413 | OR52N2 | 335 | | |
| | | | | | | | | OR51S1 | 368 | OR52N4 | 341 | | |
| | | | | | | | | OR51T1 | 23 | OR52N5 | 284 | | |
| | | | | | | | | OR51V1 | 371 | OR52R1 | 315 | | |
| | | | | | | | | | | OR52W1 | 324 | | |

**Supplementary Table S2: Number of mutations in human ORs collected in the study.** The table shows the number of nucleotide variants identified in 374 functional OR genes belonging to 17 OR families.

### Supplementary Table S3

| GPCR Domain | BW Position | Most cons. (%) hOR | Most cons. (%) Class A GPCRs | Functional Role | REFERENCE |
|---|---|---|---|---|---|
| TM1 | 1.49 | G (88%) | G (67%) | Part of the "GN" conserved motif at human and mouse ORs | PMID:26044705 |
| | 1.50 | N (99%) | N (98%) | A conserved asparagine residue occupies this position in ORs and Class A GPCRs. Involved in hydrogen bond network with D2.50 and N7.49 stabilizing the TM1, TM2 and TM7 domain region | PMID: 9115256 |
| TM2 | 2.50 | D (81%) | D (92%) | Negatively charged (D/E) residues occupy this position in most Class A GPCRs. Participates in stabilizing hydrogen bond networks with TM1 and TM7 residues. Also involved in the coordination of ions in some receptors | PMID: 29395784<br>PMID: 31855179 |
| | 2.53 | Y (43%)<br>F (23%)<br>L (20%) | V (30%)<br>F (21%)<br>M (13%) | In Class A GPCRs, this position contains >80% of bulky/aromatic residues. May be implicated in the first stage of activation pathway through TM2-TM7 | PMID: 24041646 |
| | 2.59 | P (98%) | P (36%)<br>F (21%)<br>L (18%) | A conserved proline in this position induces a structural bulge in the TM2 in several class A GPCRs | PMID: 22435816 |
| TM3 | 3.25 | C (98%) | C (86%) | Involved in a disulfide bond with ECL2 in >80% of Class A GPCRs | PMID: 21864311 |
| | 3.39 | E (81%)<br>D (13%) | S (71%)<br>G (12%) | Identified as a hot-spot position that leads to substantially higher stability for several Class A GPCRs in the inactive state. Also associated to the coordination of ions in some receptors | PMID: 28644022 |
| | 3.40 | C (37%)<br>S (22%) | I (40%)<br>V (21%)<br>L (19%) | Part of the "transmission switch" in Class A GPCRs involved in activation | PMID: 22300046 |
| | 3.49 | D (99%) | D (68%)<br>E (23%) | Part of the "[D/E]RY" motif in Class A GPCRs involved in activation | PMID: 17192495 |
| | 3.50 | R (89%) | R (97%) | Part of the "[D/E]RY" motif in Class A GPCRs involved in activation | PMID: 17192495 |
| | 3.51 | Y (80%) | Y (72%) | Part of the "[D/E]RY" motif in Class A GPCRs involved in activation | PMID: 17192495 |
| | 3.54 | I (88%) | I (54%)<br>V (36%) | A bulky hydrophobic residue involved in interactions with G-proteins in several Class A GPCRs | PMID: 24016604<br>PMID: 25205354 |
| TM4 | 4.50 | W (57%)<br>Y (17%) | W (96%) | A conserved tryptophan residue occupies this position in most class A GPCRs. | PMID: 21921973 |
| | 4.53 | G (71%) | S (45%)<br>G (32%) | A conserved glycine in this position is crucial for cell surface trafficking of model ORs. | PMID: 31974307 |
| TM5 | 5.50 | P (39%)<br>D (14%) | P (77%) | Part of the "transmission switch" in Class A GPCRs involved in activation | PMID: 22300046 |
| | 5.57 | S (98%) | C (39%)<br>L (13%) | Involved in the Class A GPCR activation pathway in several receptors | PMID: 31855179 |
| | 5.58 | Y (97%) | Y (75%) | Involved in the Class A GPCR activation pathway in several receptors | PMID: 31855179 |
| | 5.65 | I (47%)<br>V (43%) | I (46%)<br>V (16%)<br>A (15%) | Part of the hydrophobic "[I/L]xxL" motif at the intracellular end of TM5. Involved in interactions with G-proteins in several receptors. Mutations, particularly to polar amino acids, at this position in class A GPCRs inhibit G-protein coupling. | PMID: 23235263 |
| TM6 | 6.30 | R (63%)<br>K (12%) | E (35%)<br>K (15%)<br>R (14%) | Part of the "ionic lock" in Class A GPCRs involved in activation | PMID: 22300046 |
| | 6.32 | K (92%) | K (44%)<br>R (32%) | Involved in interactions with G-proteins in several Class A GPCRs | PMID: 23245528 |
| | 6.33 | A (90%) | A (31%)<br>V (19%)<br>L (10%) | Involved in interactions with G-proteins in several Class A GPCRs | PMID: 23245528 |
| | 6.37 | C (95%) | L (39%)<br>V (22%)<br>I (19%) | A highly conserved cysteine in human ORs. A bulky hydrophobic residue in this position is involved in the TM6 movement during Class A GPCRs activation | PMID: 29498889 |
| | 6.44 | V (84%) | F (80%) | Part of the "transmission switch" in Class A GPCRs involved in activation | PMID: 22300046 |
| | 6.48 | Y (69%)<br>F (23%) | W (78%)<br>F (9%) | Part of the WxP motif on TM6 in the majority of Class A GPCRs involved in activation | PMID: 22032986<br>PMID: 19375807 |
| | 6.50 | P (33%)<br>T (33%)<br>A (33%) | P (98%) | Part of the WxP motif on TM6 in the majority of Class A GPCRs involved in activation | PMID: 22032986<br>PMID: 19375807 |
| TM7 | 7.46 | P (95%) | S (60%)<br>C (13%)<br>A (13%) | A highly conserved proline in human ORs. Participates in an extended H-bond network important for receptor activation | PMID: 20395291<br>PMID: 20192770 |
| | 7.49 | N (97%) | N (77%) | Part of the NP7.50xxY motif essential for forming the active conformation, also participates in forming the G protein–binding site. | PMID: 29925258 |
| | 7.50 | P (97%) | P (96%) | Part of the NP7.50xxY motif essential for forming the active conformation, also participates in forming the G protein–binding site. | PMID: 29925258 |
| | 7.53 | Y (96%) | Y (92%) | Part of the NP7.50xxY motif essential for forming the active conformation, also participates in forming the G protein–binding site. | PMID: 29925258 |
| ECL2 | 45.50 | C (99%) | C (>80%) | Involved in a disulfide bond with the extracellular side of the TM3 in the majority of class A GPCRs | PMID: 21864311 |

**Supplementary Table S3: Conserved topological sites with functional implication in the GPCR activity.** The table shows the topological domain location, BW number, type and percentages of most conserved amino acids in 30 topological positions identified as important for the function of class A GPCRs according to several studies. Conservation values were taken from the MSA of Supplementary Figure 1 and from (http://lmc.uab.cat/gmos/).

### Supplementary Table S4

| UniProtKB entry name | Receptor Name | Organism | Resolution (Å) | Ligand Name | Ligand Function | PDBid |
|---|---|---|---|---|---|---|
| **5HT2A** | 5-Hydroxytryptamine receptor 2A | Human | 2.9 | Zotepine | Antagonist | 6A94 |
| **5HT1B** | 5-Hydroxytryptamine receptor 1B | Human | 2.8 | Dihydroergotamine | Agonist | 4IAQ |
| **5HT2B** | 5-Hydroxytryptamine receptor 2B | Human | 2.7 | Ergotamine | Agonist | 4IB4 |
| **5HT2C** | 5-Hydroxytryptamine receptor 2C | Human | 2.7 | Ritanserin | Inverse agonist | 6BQH |
| **AA1R** | Adenosine receptor A1 | Human | 3.2 | CHEMBL144360 | Antagonist | 5UEN |
| **AA2AR** | Adenosine receptor A2a | Human | 2.7 | ZM241385 | Antagonist | 3VG9 |
| **ACM1** | Muscarinic acetylcholine receptor M1 | Human | 2.7 | CHEMBL258622 | Antagonist | 5CXV |
| **ACM2** | Muscarinic acetylcholine receptor M2 | Human | 2.3 | N-methyl scopolamine | Antagonist | 5ZKC |
| **ACM4** | Muscarinic acetylcholine receptor M4 | Human | 2.6 | Tiotropium | Antagonist | 5DSG |
| **ADRB1** | Beta-1 adrenergic receptor | Turkey | 2.3 | (S)-Carvedilol | Inverse agonist | 4AMJ |
| **ADRB2** | Beta-2 adrenergic receptor | Human | 3.2 | Carazolol | Inverse agonist | 5JQH |
| **AGTR1** | Type-1 angiotensin II receptor | Human | 2.8 | OLM | Inverse agonist | 4ZUD |
| **CNR1** | Cannabinoid receptor 1 | Human | 2.8 | SCHEMBL662960 | Antagonist | 5TGZ |
| **CNR2** | Cannabinoid receptor 2 | Human | 2.8 | AM10257 | Antagonist | 5ZTY |
| **CXCR4** | C-X-C chemokine receptor type 4 | Human | 2.5 | IT1t | Antagonist | 3ODU |
| **DRD2** | Dopamine D2 receptor | Human | 2.9 | Risperidone | Inverse agonist | 6CM4 |
| **DRD3** | Dopamine D3 receptor | Human | 2.9 | Eticlopride | Antagonist | 3PBL |
| **DRD4** | Dopamine D4 receptor | Human | 2.1 | Nemonapride | Antagonist | 5WIV |
| **EDNRB** | Endothelin receptor type B | Human | 2.2 | K-8794 | Antagonist | 5X93 |
| **HRH1** | Histamine H1 receptor | Human | 3.1 | Doxepin | Antagonist | 3RZE |
| **MTR1A** | Melatonin receptor type 1A | Human | 2.8 | Ramelteon | Agonist | 6ME2 |
| **NK1R** | Tachykinin receptor 1 | Human | 2.2 | Netupitant | Antagonist | 6HLP |
| **NPY1R** | Neuropeptide Y receptor type 1 | Human | 3.0 | BMS-193885 | Antagonist | 5ZBH |
| **OPRD** | Delta-type opioid receptor | Human | 1.8 | Naltrindole | Antagonist | 4N6H |
| **OPRK** | Kappa-type opioid receptor | Human | 2.9 | JDTic | Antagonist | 4DJH |
| **OPRM** | Mu-type opioid receptor | Mouse | 2.8 | BF0 | Antagonist | 4DKL |
| **OPRX** | Nociceptin receptor | Human | 3.0 | N/A | N/A | 5DHG |
| **OPSD** | Rhodopsin | Bovine | 2.7 | Retinal | Inverse agonist | 1GZM |
| **OX1R** | Orexin receptor type 1 | Human | 2.8 | Suvorexant | Antagonist | 4ZJ8 |
| **OX2R** | Orexin receptor type 2 | Human | 2.3 | EMPA | Antagonist | 5WS3 |
| **P2RY1** | P2Y purinoceptor 1 | Human | 2.7 | MRS2500 | Antagonist | 4XNW |
| **P2Y12** | P2Y purinoceptor 12 | Human | 2.6 | AZD1283 | Antagonist | 4NTJ |
| **PAR1** | Proteinase-activated receptor 1 | Human | 2.2 | Vorapaxar | Antagonist | 3VW7 |
| **PTAFR** | Platelet-activating factor receptor | Human | 2.9 | ABT-491 | Inverse agonist | 5ZKQ |
| **LPAR1** | Lysophosphatidic acid receptor 1 | Human | 2.9 | ONO-9910539 | Antagonist | 4Z35 |
| **S1PR1** | Lysophospholipid (S1P) | Human | 2.8 | 909725-61-7 | Antagonist | 3V2Y |
| **PE2R3** | Prostanoid | Human | 2.5 | Misoprostol-FA | Antagonist | 6M9T |
| **PE2R4** | Prostanoid | Human | 3.2 | ONO-AE3-208 | Antagonist | 5YWY |
| **TA2R** | Prostanoid | Human | 3.0 | Daltroban | Antagonist | 6IIV |

**Supplementary Table S4: Non-olfactory class A GPCRs used in topological annotation.** The table shows information of non-olfactory receptors with solved 3D-structures used in topological annotation and ligand binding site (BS) definition of OR natural variants (see Supplementary Figures S2 and S4).

**Supplementary Table S5: The human OR mutation database table.** The mutation data table is available at http://lmc.uab.cat/hORMdb and contains information of 118,057 natural human OR nucleotide variants extracted from gnomAD (https://gnomad.broadinstitute.org/) and annotated with genomic and structural information as described in the main text (resumed in the diagram of Figure 1). A total of 78 descriptors were associated to each natural variant generating a total of 9,208,446 data points. More information about the data types at each column can be found at the HELP panel on the web server application.
